# Plasmin-mediated cleavage of EphA4 at central amygdala inhibitory synapses controls anxiety

**DOI:** 10.1101/2021.07.16.452595

**Authors:** Mariusz Mucha, Alberto Labrador-Ramos, Benjamin K. Attwood, Malgorzata Bajor, Jaison B. Kolenchery, Anna E. Skrzypiec, Valentina Brambilla, Marta Magnowska, Izabela Figiel, Michal Swiatek, Lucja Wiktorowska, Rahul S. Shah, Barbara Pijet, Yusuke Sakai, Nobuo Nagai, Agata Klejman, Jakub Wlodarczyk, Leszek Kaczmarek, Ryszard Przewlocki, Robert Pawlak

## Abstract

Severe stress can trigger complex behavioural changes such as high anxiety (1). Inhibitory GABA-ergic interneurons in the lateral division of the central amygdala (CEl) control anxiety through feedforward inhibition of their target cells in the medial division (CEm) (2, 3). In particular, PKCδ-positive (PKCδ^+^) interneurons in CEl are critical elements of the neuronal circuitry of fear and anxiety (3–5), but the molecular mechanisms they employ are poorly understood. Here, we show that, during stress, GABA-ergic synapses of amygdala PKCδ^+^ interneurons are regulated by a serine protease plasmin. On stress, plasmin cleaves the extracellular portion of the tyrosine kinase receptor EphA4 triggering its dissociation from gephyrin, a postsynaptic GABA-receptor anchoring protein. Dynamic EphA4/gephyrin interaction leads to modification of dendritic spine morphology and synaptic GABA-receptor expression profile. Consistent with the critical role for the plasmin/EphA4/gephyrin signalling axis in anxiogenesis, viral delivery of plasmin-resistant (prEphA4) form of EphA4 into the central amygdala prevents the development of stress-induced anxiety in mice, while the delivery of plasmin-truncated EphA4 (tEphA4) dramatically enhances this effect. Thus, our studies identify a novel, critical molecular cascade regulating GABA-ergic signalling in the central amygdala synapses that allows bidirectional switching of animal behaviour from high to low anxiety states.

## Introduction

Stress triggers protective physiological responses that are directed toward maintaining homeostasis and promoting balanced, adequate behavioral reactions. However, severe or sustained stress can compromise this response and lead to the development of neuro-behavioural abnormalities, such as cognitive impairments, depression and elevated levels of anxiety (1, 6). Anxiety disorders (such as panic disorder, specific phobias, obsessive-compulsive disorder, generalized anxiety disorder or the post-traumatic stress disorder) collectively constitute the most commonly diagnosed group of psychiatric conditions, with a life-time prevalence of ~30% (7). As such, anxiety disorders generate an enormous economic and social impact that cannot be efficiently tackled due to low efficiency of available anxiolytic therapies (7, 8). Lack of success in developing effective anxiolytics is a consequence of superficial understanding of the topography of anxiety circuits and insufficient knowledge of synaptic events underpinning stress-induced behavioural changes.

The amygdala is critically involved in the development of anxiety (1, 3, 9). Although significant progress has been made in understanding of the amygdala’s role and wiring design, full comprehension of its function requires deciphering how its cellular architecture, neuronal projections and synaptic machinery work in concert to control behaviour.

The amygdala is composed of several groups of nuclei, including the basolateral (BLA; composed of the lateral nucleus LA, and basal nucleus BA); central (CeA, further divided into the lateral Cel, and medial CeM), medial amygdala (MeA) and intercalated cell masses (ITCs) (3, 9, 10). The amygdala is composed of a variety of cell types, with diverse genetic and biochemical profiles (11). Overall, the amygdala sub-regions utilize either excitatory glutamatergic transmission (neurons located primarily in the BLA) or inhibitory GABA-ergic signaling (interneurons that predominantly form CeA, MeA, ITCs) (3, 9). CeA neurons transmit emotionally significant neural signals to downstream brain areas that confer direct expression of behavioural outcomes (e.g. risk avoidance, anhedonia, anorexia, social avoidance, high blood pressure, panting, and heart palpitations), which jointly constitute the anxiety-like state (3, 9). To this end, CEl-located interneurons that express protein kinase Cδ (PKCδ) are principal gate-keepers controlling the activity of downstream projection neurons (4, 12) and have recently been implicated in controlling anxiety-like behaviour (4).

Synaptic proteolysis provides an attractive mechanism by which neuronal transmission in the amygdala could be regulated (13–16). Stimulus-evoked release of a protease to the synaptic cleft could control neuronal plasticity by cleaving local extracellular matrix proteins, shedding membrane receptors, degrading adhesion molecules and activating zymogen proteases such as plasminogen (13–16). Proteases can be released in response to neuronal depolarization, regulate multiple forms of synaptic plasticity and learning (16–18) and anxiety-like behavior (14).

Eph-receptor tyrosine kinases are an important group of synaptic plasticity-related molecules that can be cleaved by extracellular proteases (13, 19). For instance, we have recently demonstrated that neuropsin-mediated cleavage of EphB2 controls excitatory transmission in the BLA synapses and promotes stress-induced anxiety by regulating the interaction between EphB2 and the NR1 subunit of the NMDA receptor (13). Although analogous, protease-mediated mechanisms controlling inhibitory transmission in the CeA would be attractive, such a hypothesis has never been proposed or examined.

Here we identify a novel signalling cascade controlling inhibitory plasticity at the CEl synapses that allows bidirectional switching of animal behaviour to high or low anxiety states. On stress, CEl PKC^+^ interneurons release tPA to convert an inactive zymogen plasminogen to active protease plasmin. In turn, plasmin cleaves ephrin receptor EphA4 triggering its dissociation from a master regulator of GABA-receptor anchoring, gephyrin. Dynamic EphA4/gephyrin interaction promotes changes in the amygdala GABA-receptor expression profile and in dendritic spine morphology. Consistently, overexpression of the plasmin-resistant form of EphA4 (prEphA4) in the central amygdala prevents, while overexpression of plasmin-truncated form of EphA4 (tEphA4) enhances, stress-induced anxiety in mice.

## Material and Methods

### Mice

Experiments were performed on 8-12-week-old male wild-type (C57BL/6J), tPA^−/−^ or plasminogen^−/−^ mice backcrossed to C57BL/6J for 12 generations. All mice were housed in groups of three to five male mice per cage in a colony room with a 12 hour light/dark cycle in standard cages with ad libitum access to commercial food pellets and water. All experiments were conducted during the light half of the cycle. The experiments were approved by the appropriate national and local ethics committees.

### Amygdala dissection

Mice (stress-naive or subjected to the restraint stress as described below) were euthanized using intraperitoneal sodium pentobarbital (50 mg/kg) and perfused transcardially with ice-cold PBS. The brains were removed and amygdalae dissected from a slice, −0.58 to −2.3mm relative to Bregma using a brain matrix (Stoelting), frozen immediately on dry ice and stored at −80°C until use.

### Microarray study

Amygdalae were isolated from tPA+/+ (wild-type; n = 30) and tPA−/− (n = 30) mice using a dissecting microscope in ice-cold ACSF (glucose 25 mM, NaCl 115 mM, NaH2PO4·H2O 1.2 mM, KCl 3.3 mM, CaCl2, 2 mM, MgSO4, 1 mM, NaHCO3 25.5 mM, pH 7.4 and stored at −20 °C in RNAlater solution (Qiagen). RNA was extracted using RNeasy Lipid Tissue Mini Kit (Qiagen), the ribosomal fraction of RNA reduced with RiboMinus Kit (Invitrogen) and the RNA integrity verified by electrophoresis using Agilent Bioanalyser 2100 (Agilent Technologies). RNA pulled from three mice was reverse-transcribed and hybridized with GeneChip Mouse Exon 1.0 ST Array (Affymetrix; 10 arrays per genotype).

The Bioconductor bundle of R packages and The Partek Genomics Suite (PGS) have been employed to analyse differential gene expression and perform pathway analyses. The results were verified using Enrichr and g:Profiler servers using KEGG Pathway Database (www.kegg.jp) as the basis for performing Statistical Overrepresentation Test.

### Cell culture experiments

SHSY-5Y cells were incubated (37°C, 5% CO_2_) in medium (MEM + Earle’s Balanced Salt Solution, 5% fetal calf serum, 5% new-born calf serum, 2 mM L-glutamine, 1% penicilin-streptomycin) until 80-90% confluence. They were washed with PBS (+Ca^2+^, +Mg^2+^) three times before being incubated with PBS, PBS + tPA (Alteplase, Genentech; 1 µg/ml) or PBS + tPA (Alteplase, Genentech; 14 nM) + plasminogen (R&D, 6, 12 or 120 nM) for 15 min, after which the dishes were placed on ice and protease inhibitors (cOmplete^®^, Roche) were added. The cells were collected using a cell scraper and homogenised (Tris HCl pH 7.4 50 mM, NaCl 150 mM, EDTA 5 mM, EGTA 5mM, Triton-100 1%, NP40 0.5%, pH7.5). The resulting protein sample was analysed by Western blotting as described below.

Neuro-2A cells were incubated (37°C, 5% CO_2_) in cell culture media (DMEM, 1% penicilin-streptomycin, 1% non-essential amino-acids) + 5% v/v fetal bovine serum (FBS), until 70-80% confluence. Cells were washed twice with culture medium without FBS before being incubated with culture media without FBS for 3 h. Then, cells were treated with medium + t-PA (Invitrogen, USA; 14 nM) or t-PA + plasminogen (R&D Systems, UK; 6, 12 or 120 nM) for 15 min. After tPA/plasminogen incubation, culture dishes were placed on ice, rinsed twice with ice-cold phosphate buffer saline, pH=7.4 (PBS) and cells were lysed in RIPA buffer (50 mM Tris HCl pH 7.4, 150 mM NaCl, 1% NP-40, 0.5 % deoxycholate, 0.1% sodium dodecyl sulfate (SDS), 1 mM ethylenediaminetetraacetic acid (EDTA) solution, 10 mM NaF, 1 mM sodium vanadate, 1x Inihbitor Cocktail (cOmplete^®^, Roche, 1x protease inhibitors (Halt^®^, Thermo)) followed by homogenization with a 25 gauge needle and a syringe on ice. The resulting protein lysates were analysed by Western blotting as described below.

For imaging experiments, coverslips were flame sterilized, polylysine coated and put on culture plates. Neuro-2A cells were cultured in culture medium + 5% FBS and transfected with a plasmid expressing EphA4 variants using polyethylenimine (PEI). The backbone plasmid used to transfect cell lines was pIRES2-EGFP2 (Clontech, UK). Coverslips were fixed with 4% paraformaldehyde (PFA) in PBS and washed 3 times for 15 min with PBS and treated with PBS + 0.1% Triton X-100 + 10% FBS for 1 h at room temperature (RT). Samples were then probed with the following primary antibodies: mouse anti-EphA4, (Invitrogen, USA, 1:500), anti-GFP antibody (Santa Cruz Biotechnology Inc., USA, 1:500) at 4°C overnight with gentle agitation. The next day, samples were washed (PBS + 0.1% Triton X-100 for 3×15 minutes at RT) before applying the corresponding secondary fluorescent antibodies (in PBS + 0.1% Triton X-100 + 10% FBS for 1 h at RT in darkness; Invitrogen, USA). Images were taken with Zeiss LSM5 Exciter and processed with Zen 2009 (Zeiss Ltd., Germany).

### Immunohistochemistry

Mice were euthanized and perfused transcardially with ice-cold PBS followed by 4% PFA in PBS. The brains were extracted and fixed in cold 4% PFA in PBS overnight. The next day, brains were washed in PBS. Then, coronal brain sections of 70 µm were cut using a vibrating microtome (Campden Instruments, UK). Sections were incubated for 20 minutes at RT in PBS + 3% H_2_O_2_ with gentle rotation and then washed 3×10 min with PBS. After treatment with PBS + 0.1% Triton X-100 + 10% FBS (for 1 h at RT), slices were probed with the primary antibodies at 4°C overnight with gentle agitation. The following antibodies were used: mouse anti-EphA4 (Invitrogen, USA; 1:500); rabbit anti-plasminogen (Innovative Research, USA; 1:500); rabbit anti-tPA (Molecular Innovations, UK; 1:500), mouse anti-NeuN (Chemicon, 1:1000, chicken anti-gephyrin (Abcam, USA; 1:500). Then, the sections were washed (PBS + 0.1% Triton X-100 for 3×15 minutes at RT) before applying the corresponding secondary fluorescent antibodies (Invitrogen, USA; 1:1000) (in PBS + 0.1% Triton X-100 + 10% FBS for 1 h at RT in darkness).

For immunostaining with amplified fluorescence, secondary fluorescent antibodies were substituted with horse anti-rabbit biotin-conjugated antibodies (Vector Labs, UK, 1:500). After incubation, slices were washed for 3×15 minutes with 1x PBS + 0.1% Triton X-100 followed by 45 minutes incubation at room temperature with ABC reagent (Vector Laboratories) and then Tyramide Signal Amplification (TSA) Systems kit (Perkin Elmer) was used according to manufacturer instructions. Next, slices were washed 3×15 minutes with PBS + 0.1% Triton X-100 before mounting.

### In situ hybridization

682bp long fragment of mouse tissue plasminogen activator mRNA 3’UTR was amplified by Taq PCR with 5’-GTGCCTGGGGTCTACACAAA and 5’-AAATCATACAGTTCTCCCAACCA primers from mouse amygdala cDNA. The PCR product was next cloned into a dual promoter pCRII vector and the insert orientation and integrity was confirmed by restriction analysis and DNA sequencing. The plasmid was linearized with SacI or XhoI (for transcription from T7 or SP6 promoter respectively) and 3’UTR fragment was transcribed *in vitro* using SP6 or T7 polymerases (New England Biolabs, UK) and the DIG RNA labelling Mix (Roche), generating sense and antisense DIG labelled probes.

Mouse brains were fixed overnight in 4% PFA in PBS at 4°C. The next day, brains were washed in PBS treated with DEPC and sectioned at 50 µm. The sections were transferred onto polylysine slides (VWR, UK), left to dry and stored at −80 degrees until use. Before *in situ* hybridisation commenced, slides were submerged in PBS/DEPC until the PBS precipitate dissolved. In the meantime the RNA probe was incubated with hybridisation buffer (50% v/v deionized formamide, 0.2M NaCl, 50mM EDTA, 10mM Tris-HCl, pH7.5, 5mM NaH_2_PO_4_ · 2H_2_O, 5mM Na_2_HPO_4_, 0.05 mg/ml tRNA from baker’s yeast) for 5 minutes at 70°C and placed onto slides. The dilution of probe in the hybridisation buffer generating the best signal to noise ratio was established empirically as 1:1000. Sections were covered with cover glasses and the hybridization was performed at 65°C overnight in a chamber humidified with 50% v/v formamide containing 1× SSC buffer. The next day sections were washed three times at 65°C for 30 min each in wash solution (50% v/v formamide, 1× SSC, 0.1% Tween 20), followed by two washes for 30 minutes each in 1× MABT (100 mM maleic acid, 150 mM NaCl, pH 7.5, 0.1% Tween 20) at RT. The sections were blocked with blocking buffer (0.5% blocking reagent (Roche) dissolved in TNT buffer (0.1M Tris-HCl pH7.5, 0.15M NaCl, 0.05% Tween-20)) for 1 hour at RT. Anti-Digoxigenin-POD (Roche) antibody at 1:200 dilution in blocking buffer was applied onto the sections and incubated for 1 hour at RT, followed by 3 washes with PBST (PBS containing 0.1% Triton X-100). The biotin deposition was performed using TSA Plus Biotin Kit (Perkin Elmer), followed by 3 subsequent washes with PBST and 45 minute incubation with ABC reagent (Vector Laboratories). Then, fluorescein was deposited using TSA Plus Fluorescein Kit (Perkin Elmer). To immunohistochemically visualise CRH, SOM and PKCδ proteins on the same sections, the slides were blocked with 10% FBS in PBST for 1 hour at RT followed by overnight incubation at 4°C with either anti-CRH (Sigma, UK), anti-SOM (Merck Millipore, UK) or anti-PKC-delta (BD Biosciences, UK) primary antibodies diluted 1:200. The primary antibodies were visualised by corresponding AlexaFluor-conjugated secondary antibodies (Invitrogen, USA) as described in the Immunohistochemistry section.

### Eph receptor cleavage

Recombinant Mouse EphA4-Fc Chimera Protein, CF (R&D Systems, UK; 1 µg/ml) was incubated with t-PA (14 nM) or t-PA + plasminogen (6, 12 or 120 nM) or without proteases in a HEPES-Tween buffer (0.1 M HEPES, 0.01% Tween, pH 7.4) for 15 min. The samples were placed on ice and protease inhibitor cocktail (cOmplete^®^, Roche) was added to stop the reaction. The samples were then analysed by SDS-PAGE and Western blotting as described below.

### EphA4 cleavage in brain homogenates

Amygdalae were extracted and homogenized in 0.1 M Tris, 0.1% Triton X-100, pH 7.4, containing phosphatase inhibitors (10 mM NaF, 1 mM Na_3_VO_4_). The homogenate (100 µl) was incubated with t-PA (Invitrogen, USA; 14 nM) alone, t-PA + plasminogen (R&D Systems, UK; 6, 12 or 120 nM) or without proteases for 15 minutes at 37°C. The homogenate was placed on ice and protease inhibitor cocktail was added to stop the reaction. The samples were then analysed by SDS-PAGE and Western blotting as described below.

### Western blotting

Samples were homogenized in 0.1 M Tris, 0.1% Triton X-100, pH 7.4, containing phosphatase inhibitors (10 mM NaF, 1 mM Na_3_VO_4_) and protease inhibitors (cOmplete^®^, Roche) and the protein concentration was adjusted to 2 mg/ml using the Bradford method (Thermo Scientific, USA). Samples were then reduced using DTT (Sigma-Aldrich, USA), denatured (100°C for 5 min), subjected to SDS-PAGE electrophoresis and transferred onto a nitrocellulose membrane (Thermo Scientific, USA). After blocking (5% skim milk in Tris-buffered saline + 0.01 % Tween^®^20 (Sigma-Aldrich, USA) for 1 h at RT), the membranes were probed with the following primary antibodies at 4°C overnight: mouse anti-EphA4 C-terminus (Invitrogen, USA; 1:1000), goat anti-EphA4 N-terminus (R&D Systems, UK; 1:500), goat anti-EphB2, goat anti-EphB6 and goat anti-EphrinB2 (R&D, 1:500, 1:500 and 1:300 respectively) and rabbit anti-p75NGF receptor (Chemicon, USA, 1:1000). The membranes were then washed in PBST (3×5 minutes) before incubation with a relevant HRP-conjugated secondary antibody as appropriate (Vector Labs, UK; 1:1000, 1 h, RT). The signal was developed, after washing with PBST (3×15 min), using Amersham ECL detection reagent (GE Healthcare, USA). To normalize the results, all membranes were stripped using a Restore PLUS buffer (Pierce, UK), blocked, washed as above described and re-blotted using mouse anti-β-actin antibody (Sigma-Aldrich, USA; 1:2500, 1 h, RT). Again, the membranes were washed, incubated and developed as described above. To quantify the results, the band intensities were normalized to β-actin levels.

### Cleavage of EphA4-Fc for ATOMS

Lyophilized EphA4Fc (R&D, #641-A4), plasminogen (R&D #1939-SE) or tPA (Alteplase) were reconstituted in 500µl 50 mM HEPES buffer, pH 7.4. To remove the interfering compounds contained in the protein solution, 50 mM HEPES buffer, pH 7.4 was exchanged three times using Millipore Amicon Ultra devices with cut off 3000 Da (#UFC500324) at 10 000 x g for 10 minutes at 4°C. The filtrates from the three steps were pooled and the total amount of protein in all samples was determined by UV absorption A_280_. EphA4Fc (10µg/ml) was incubated either with tPA (14 nM) or tPA + plasminogen (12, 120 or 240 nM) or without proteases in a 50 mM HEPES buffer, pH 7.4 for 15 min at 37°C. tPA + plasminogen were pre-incubated for 15 min at 37°C before mixing with EphA4Fc. Then, cleavage reaction samples were subjected to the ATOMS analysis.

### ATOMS analysis

To each sample one volume of 8.0 M GuHCl was added to denature proteins. The pH of the samples was adjusted to 7.0 and DTT (to final concentration 5 mM) was added to reduce disulfide bridges. Samples were incubated at 65°C for 1 h. Then, iodoacetamide (to final concentration of 15 mM) followed by DTT (to final concentration of 30 mM) were added and samples were incubated at RT in the dark for 30 min and at RT for 30 min, respectively. Then samples were labeled with either heavy formaldehyde (formaldehyde containing the isotope ^13^C and deuterium (^13^CD_2_O from Cambridge Isotope Laboratories, Inc) to a final concentration of 60 mM or light formaldehyde (regular formaldehyde (^12^CH_2_O from Sigma) to a final concentration of 60 mM. Next, NaBH_3_CN to a final concentration of 30 mM was added to all samples. Samples were vortexed and the pH was adjusted to 6-7 followed by overnight incubation at 37°C. To quench the excess formaldehyde, ammonium bicarbonate (final concentration, 100 mM) was added and pH was adjusted once again to 6-7. Samples were incubated at 37°C for 4 h. Then, samples were combined as appropriate and precipitated with cold acetone/methanol. After precipitation, dried protein pellets were re-suspended with 60 µl of 50 mM HEPES, pH 8.0 and 1 µg of mass spectrometry grade trypsin was added to each sample followed by overnight incubation at 37°C. Then samples were subjected to mass spectrometry analysis.

### Mass spectrometry analysis

The resulting peptide mixtures were applied to an RP-18 pre-column (Waters, Milford, MA, USA) using water that contained 0.1% formic acid as a mobile phase and then transferred to the RP-18 column (75 µM internal diameter; Waters) of the nanoACQUITY UPLC system (Waters) using an ACN gradient (0–30% ACN in 45 min) in the presence of 0.1% formic acid at a flow rate of 250 nL/min. The column outlet was coupled directly to the ion source of an LTQ Orbitrap Velos mass spectrometer (Thermo Electron, San Jose, CA, USA) working in the regime of data-dependent MS to MS/MS switch. A blank run that ensured the absence of cross-contamination from previous samples preceded each analysis. The obtained mass spectra were preprocessed with Mascot Distiller software (v. 2.2.1, Matrix Science) and searched against the EphA4-Fc protein sequence using on-site-licensed-processor-engine MASCOT software (Mascot Server v. 2.2.03, Mascot Daemon v. 2.2.2, Matrix Science). Fixed modification for carboxymethylation of cysteines and variable modification of methionine oxidation were applied to all searches. Enzyme specificity was semi-Arg-C, precursor and fragment ion mass tolerance was 0.8 Da, peptide mass tolerance was 40 ppm, and a maximum of three miscleavages were allowed. The variable modifications including lysine and N-terminal dimethylation with heavy formaldehyde (34.0631 Da) and with light formaldehyde (28.0311 Da) were conducted for all samples. Protein MASCOT scores above expectation values of 0.05 were required for a hit.

### EphA4 construct generation

For the generation of prEphA4 variant mutation, single amino acid substitution R497Q was performed by site directed mutagenesis assay. PCR reaction was performed using 20 ng of plasmid carrying cDNA of wild type mouse EphA4 as a template and the pair of self-reverse-complement primers introducing the required mutation (5’-GTTTTTCACGTGCGAGCCCAGACCGCTGCTGGCTACGG-3’; 5’-CCGTAGCCAGCAGCGGTCTGGGCTCGCACGTGAAAAAC-3’) followed by the treatment with DpnI restrictase and electroporation into 5-alpha electrocompetent E. coli (New England BioLabs, UK). The introduction of desired mutation and sequence integrity was verified by DNA sequencing.

For the generation of the wtEphA4, prEphA4 and tEphA4 expressing lentiviral constructs, their respective cDNA were amplified by PCR from wild type mouse EphA4 cDNA or mutated prEphA4 cDNA using the following primers (wtEphA4 and prEphA4: 5’-AAA<u>TCTAGA</u>ATGGCTGGGATTTTCTATTTC-3’ and 5’-AAA<u>GTCGAC</u>TCAGACAGGAACCATCCTGCC-3’ containing respectively XbaI and SalI restriction sites (underlined) allowing their further cloning into lentiviral vectors. The truncated form of EphA4 (tEphA4) was amplified using 5’-AAA<u>GCTAGC</u>**ATGGCCGGCATCTTCTACTTCATCCTGTTCTCCTTCCTGTTCGGCATCTG CGACGCC**ACCGCTGCTGGCTACGGAGACTTCAGC-3’ containing a NheI site and wtEphA4 signal peptide allowing tEphA4 membrane localisation (bold) and the reverse primer containing SalI site mentioned above. Sequence integrity was verified by DNA sequencing. For overexpression of EphA4 receptor and its variants, these inserts were subcloned into a plasmid empty backbone as described below.

### Lentiviral gene transfer system

EphA4 (or its mutated/truncated forms) was inserted into a LV-pUltra plasmid empty backbone (Addgene plasmid catalogue #24129) using NheI (restriction site at the 5′ terminus) and the SacI (restriction site at the 3′ terminus) downstream of an enhanced green fluorescence protein (EGFP), separated by a self-cleaving peptide P2A. LV-pULTRA plasmid uses an UbC (ubiquitin C) promoter.

For lentiviral production, HEK293T cells were grown in DMEM media (Sigma-Aldrich, USA D6046) supplemented with 1% penicilin-streptomicine and 10% v/v FBS until reaching a confluence of 70-80%. Cells were transfected with the addressing and packaging plasmids (pCMV delta R8.2, Addgene #12263; pCMV-VSV-G, Addgene #8454). After 48 hours of expression, the virus concentration was performed by ultracentrifugation. Infectivity was assessed with a focus forming assay (FFA). Viral particles were diluted in 2 ml of cell culture media and applied to Neuro-2a cell line monolayer in a 6 well plate at 100% confluency. EGFP-positive infective units were counted and the batches of viral particles diluted to achieve functional infectivity of 2 × 10^8^ transducing units (TU) / ml.

### Co-immunoprecipitation

Unless otherwise stated the experiments were conducted at 0-4°C. For EphA4 immunoprecipitation, frozen tissue (−80°C) was homogenised in the following buffer: 50 mM HEPES-NaOH, 1% v/v Triton X-100, 150 mM NaCl, 1 mM EGTA, 1.5 mM MgCl_2_, 1.5 mM, 10% glycerol, pH 7.4, containing phosphatase inhibitors (10mM NaF, 1mM Na_3_VO_4_) and protease inhibitors (cOmplete^®^, Roche). Protein concentration was adjusted to 10 mg/ml by the Bradford method (Pierce) and 5 mg of protein was used for each immunoprecipitation. Homogenates were incubated for 1 hour with 2 µg (1:250) of either an irrelevant IgG from the same species (Cell Signalling Technology, USA) or rabbit anti-EphA4 polyclonal antibody (Proteintech, UK). Then 50 µl of pre-washed (3×5 minutes in homogenising buffer) and equilibrated beads (Protein G Sepharose 4 Fast Flow, GE Healthcare, Sweden) were added and the sample mix was incubated overnight with gentle agitation. The next day, the sample was briefly centrifuged to pellet down the beads and the supernatant was collected for further analysis. Pelleted beads were gently resuspended in homogenising buffer 50:50 and left for 5 min with gentle agitation to wash. The washing process was repeated 10 times. The final pellet was resuspended in Laemlli buffer (4% SDS, 20% glycerol, 10% 2-mercaptoethanol, 0.004% bromophenol blue and 0.125 M Tris-HCl, pH 6.8 approx.) 50:50 and boiled for 5 min to elute proteins from the beads. Beads were pelleted down and the supernatant was analysed with SDS-PAGE and Western blotting as described above.

For gephyrin/EphA4 co-immunoprecipitation Neuro-2a cells were transfected with the above described EphA4-expressing vectors (wtEphA4, prEphA4, tEphA4 or EGFP-expressing control) and were homogenised in TBS pH 7.4, 1% NP-40 containing phosphatase inhibitors (10mM NaF, 1mM Na_3_VO_4_) and protease inhibitors (cOmplete^®^, Roche). Protein concentration was adjusted to 10 mg/ml by the Bradford method (Pierce, UK) and 5 mg of total protein was used for each immunoprecipitation. Streptavidin beads (Pierce(tm), ThermoFisher Scientific, USA) were pre-cleared with BSA 5% and homogenising buffer. Homogenates were incubated in 5% BSA with 1 µg (1:400) of either a biotynylated anti-gephyrin IgG (Synaptic Systems, Germany; #147111BT) or a biotynylated irrelevant IgG from the same species (Abcam, UK). Then, 50 µl of pre-washed (3×5 min in homogenising buffer); and equilibrated beads (Pierce(tm), ThermoFisher Scientific, USA) were added to the samples and the mix was incubated overnight with gentle agitation. The next day, the sample was centrifuged briefly to pellet down the beads and the supernatant was stored for further analysis. After that pelleted beads were gently resuspended in washing buffer containing 4xTBS, 5% NP-40, phosphatase inhibitors (10mM NaF, 1mM Na_3_VO_4_) and protease inhibitors (cOmplete^®^, Roche) and washed 10 times with gentle agitation. The final pellet was resuspended in Laemlli buffer (4% SDS, 20% glycerol, 10% 2-mercaptoethanol, 0.004% bromophenol blue and 0.125 M Tris-HCl, pH 6.8) 50:50 and boiled for 5 min to elute proteins from the beads. Beads were pelleted down and the supernatant was analysed with SDS-PAGE and Western blotting as described above.

### Amygdala cultures and dendritic spine analysis

For morphological analysis of dendritic spines, cells were transfected with Lipofectamine 2000 (Invitrogen, UK) according to the manufacturer protocol at 7-9 day *in vitro* (DIV) with plasmid carrying red fluorescent protein (RFP) under the β-actin promoter, together with the plasmid carrying the EphA4 receptor variants (EGFP as a control) under the CMV promoter. Imaging experiments were performed at 14 DIV.

Images were acquired using the ZEISS LSM 780 confocal microscope with a PL Apo 40×/1.4 NA oil immersion objective using 488 nm and 561 nm diode-pumped solid state lasers at 10% transmission with 1024×1024 pixel resolution. A series of z-stacks were acquired for each cell at 0.4 µm steps, with additional digital zoom that resulted in lateral resolution of 0.07 µm per pixel.

The images of dendrites were semi-automatically analyzed using the custom written software (SpineMagick software, patent no. WO/2013/021001). The dendritic spine shape parameters were determined i.e. length, head width and scale-free parameter, the length-to-width ratio (i.e. the length divided by the head width), which reflects the spine shape. The head width was defined as the diameter of the largest spine section while the bottom part of the spine (1/3 of the spine length adjacent to the dendrite) was excluded. Only the spines protruding in the transverse direction (contained in the single image plane) that could be clearly distinguished were selected. Only the spines belonging to the secondary dendrite were chosen; the motivation for this restriction was to eliminate possible systematic differences in spine morphologies due to the location of spines on dendrites with different ranks. We discarded from the analyses all objects (protrusions) with an area smaller than 0.2 µm owing to the limitation in resolution of typical confocal setup. The total number of spines analyzed was 1131 (13 cells) for wtEphA4, 1088 (14 cells) for prEphA4, 1173 (15 cells) for tEphA4 and 1241 (9 cells) for a control construct (containing EGFP only).

### Stereotaxic injections

Adult mice (8-9 weeks-old) were anaesthetized with isoflurane (5% and 4 l/min of oxygen) in an induction chamber and then were positioned in a mouse stereotaxic frame (Kopf Instruments, Germany) and anaesthetized with a constant rate of isoflurane 2.5% and 1 l/min of oxygen through a facemask. The central amygdala (CeA) was targeted bilaterally using the following stereotaxic coordinates: −1.5 mm antero-posterior (AP), ±3.0 mm medio-lateral (ML) and −4.45 mm dorsoventral (DV). After the initial injection the needle was gently withdrawn to −4.3 mm DV and the second injection delivered. Injection volumes were 500 nl delivered at 10 nl / min using a metal gauge needle attached to a NanoFil^©^ 10 µL syringe (World Precision Instruments). The needle was positioned at the first target site and remained there for 10 min before the beginning of the injection. After the injection, the needle stayed in the same position for 10 min before it was moved to the second delivery position. After the second injection was performed, the needle remained in the same position for another 10 min before it was withdrawn completely. Behavioural experiments were performed 4 weeks after recovery. All injection sites were verified by immunohistochemistry for the co-expressed EGFP, using an anti-GFP antibody (Santa Cruz Biotechnology Inc., USA; sc-9996).

### Stress and behavioural analyses

Prior to the experimental procedures mice were kept undisturbed for one week in their home cages for habituation. Restraint stress was performed in a separate room during the light period of the circadian cycle. Mice were held in plastic tube restrainers secured at the tail end of the restrainer with a cap, within their home cage (for 1-6 hours; the duration indicated in the respective Figures; 6 hours for mice subjected to behavioural analyses). Control animals were left undisturbed. 12 hours after the completion of the restraint stress mice were allowed to habituate to the behavioural room for at least 1 h prior to testing. The tests were performed during the light cycle between 09:00 and 14:00 hours.

The elevated-plus maze apparatus consisted of four non-transparent white plexiglas arms: two enclosed arms (50 × 10 × 30 cm) that formed a cross shape with the two open arms (50 × 10 cm) opposite each other. The maze was 55 cm above the floor and dimly illuminated. Mice were placed individually on the central platform, facing an open arm, and allowed to explore the apparatus for 5 min. Behaviour was recorded by an overhead camera. The time mice spent exploring the open and closed arms of the maze was determined automatically using AnyMaze software (Stoelting, UK). The maze was cleaned with 70% alcohol after each session to avoid any odorant cues.

### Statistical analysis

The data are expressed as mean ± standard error of the mean (SEM). Student’s t-test (when two groups were compared), analysis of variance (ANOVA) followed by Bonferroni’s Multiple Comparison post-test or Kruskal-Wallis H test followed by Dunn’s Multiple Comparison Test were used as appropriate. P values of less than 0.05 were considered significant. The ANOVA P values are reported in the figure legends, the results of the post-test are indicated by asterisks on graphs.

## Results

### tPA-activated plasmin regulates GABA-ergic synapses of PKCδ^+^ interneurons in the central amygdala

tPA is a serine protease rapidly induced by various forms of neuronal activity which undergoes presynaptic vesicular release upon depolarization (17). In the amygdala, tPA has previously been implicated in the regulation of anxiety (14, 20), but its mechanism of action is unknown. To identify the functional or anatomical unit regulated by tPA we compared gene expression patterns in tPA^+/+^ and tPA^−/−^ mouse amygdalae using microarray technology and performed functional pathway analysis of differentially expressed transcripts (Fig. 1a, b, c). The expression of thirteen GABA-related transcripts, including seven GABA-receptor subunits, were altered by the deletion of the tPA gene (enrichment p value <0.000000007; Fig. 1a, b, c), suggesting that tPA in the amygdala acts to regulate GABA-ergic synapses. Strikingly, tPA is expressed in amygdala subdivisions predominantly composed of inhibitory GABA-ergic interneurons (CEl) (21, 22) (Fig. 1d) but absent from regions where glutamatergic neurons predominate (basolateral amygdala) (22), consistent with a role in regulating the properties of inhibitory microcircuits.

**Figure 1:**
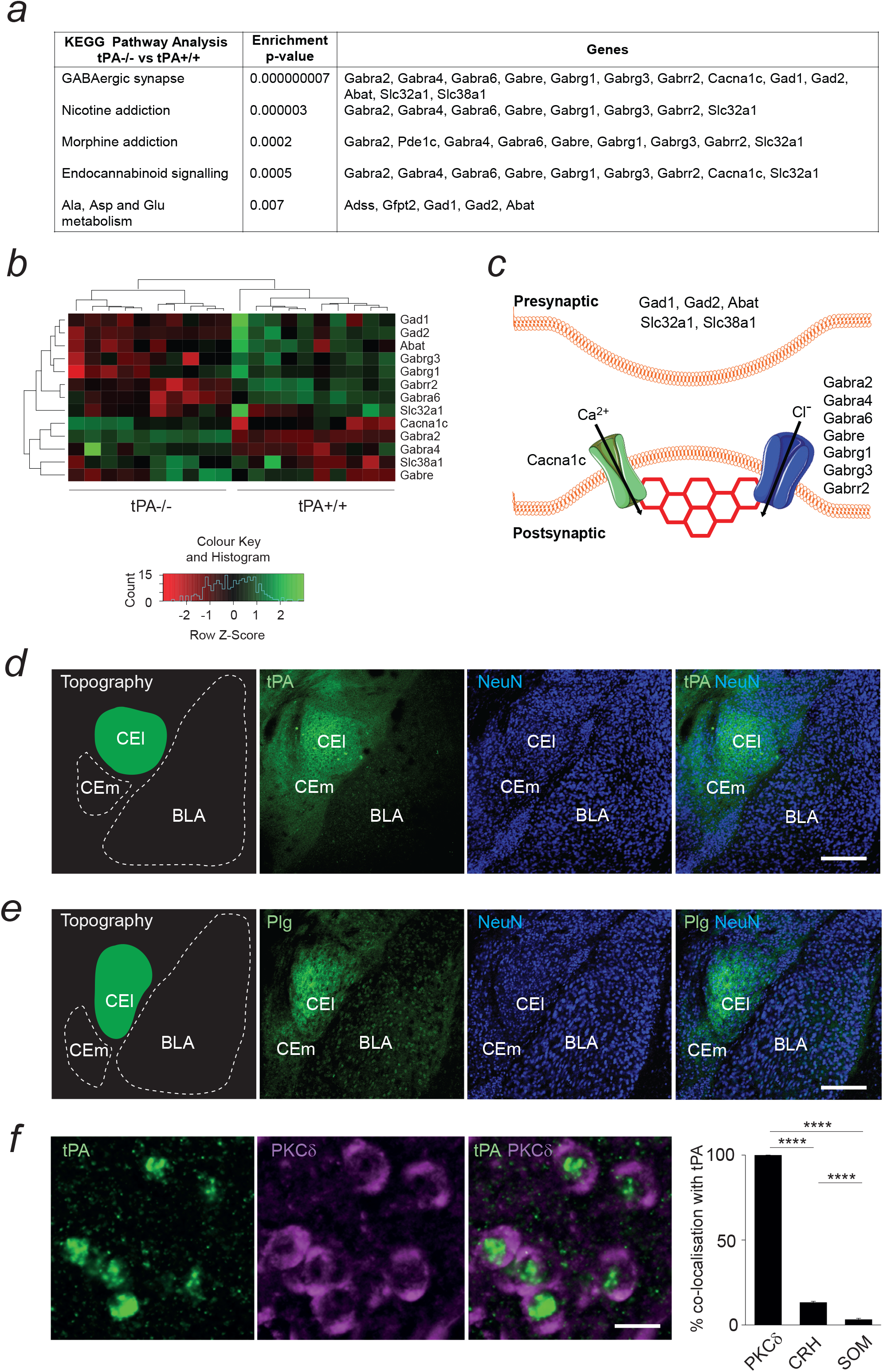
tPA is expressed by PKCd^+^ GABA-ergic interneurons in the central amygdala, co-localizes with plasminogen and regulates the composition of GABA-ergic synapses. In keeping with the role of tPA/plasminogen system in regulating GABA-ergic synapses, immunohistochemistry confirmed high levels of tPA **(d)** and plasminogen **(e)** (green) in the lateral division of the central amygdala (CEl) of the wild-type mouse where GABA-ergic transmission predominates, but not in primarily excitatory basolateral amygdala (BLA). NeuN (blue) was used to highlight cell bodies and amygdala topography. **(f)** Fluorescent in situ hybridization for the tPA mRNA (green) performed in conjunction with immunohistochemistry for protein kinase Cδ-positive (magenta), corticotropin-releasing hormone (CRH) or somatostatin (Supplementary Figure 1) revealed that tPA is mainly produced by PKCd^+^ interneurons in CEl (F_(2, 36)_ = 1903, P<0.0001). Only a small proportion of CRH^+^ or SOM^+^ cells contained tPA mRNA. **(a)** Microarray analysis of gene expression in the amygdalae of tPA^+/+^ vs tPA ^−/−^ mice followed by KEGG pathway analysis identified GABA-ergic synapses as the primary site of action of tPA in this brain region (enrichment p-value <0.000000007). Heat-map highlighting tPA-dependent changes in the GABA-ergic synapse-related transcripts is shown in **(b)**. Localization of proteins encoded by tPA-modulated genes in the GABA-ergic amygdala synapse **(c)**. Gephyrin is shown as a hexagonal postsynaptic scaffold. BLA-basolateral amygdala, CEl-lateral division of the central amygdala, CEm-medial division of the central amygdala, tPA-tissue plasminogen activator. Bars in a-b and c indicate 400 and 10 µm, respectively. Results are shown as mean ± SEM. ****p<0.0001

The extracellular pattern of tPA expression in the central amygdala (Fig. 1d) suggests either local synthesis by one or more CEl interneuron subtypes, or trafficking by long range axonal projections from other brain regions. To test the possibility of local tPA synthesis and identify the source, we combined fluorescent *in situ* hybridization for the tPA gene (*Plat*) with immunohistochemistry for cellular markers of the major interneuron subclasses populating CEl (4, 5, 21). We found that all tPA-positive neurons in the CEl co-expressed PKCδ (100±0%, n=1094 cells, N=11 sections from 3 mice), while a significantly smaller proportion expressed corticotropin-releasing factor (CRF, 13.4±0.5%, n=832 cells, N=11 sections from 3 mice, p<0.0001) or somatostatin (SOM, 3.3±0.6%, n=1526 cells, N=18 sections from 3 mice, p<0.0001)(Fig. 1f, Supplementary Fig. 1). This result points to PKCδ^+^ GABAergic interneurons, recently identified as critical elements of the neurocircuitry of anxiety (4), as being the main source of tPA in the CEl.

tPA has a narrow substrate specificity and only a few known targets (17). However, it rapidly converts the proenzyme plasminogen to the active protease plasmin, which can serve as an ultimate effector of this proteolytic cascade (17, 18). Immunohistochemistry revealed that plasminogen expression pattern in the amygdala closely resembles that of tPA, confirming the co-localization of both principal elements of the tPA/plasmin system in the CEl (Fig. 1d, e). This result, together with the outcome of the microarray study (Fig. 1a, b), suggests that synthesis and activity-dependent synaptic release of tPA by PKCδ^+^ interneurons may generate local plasmin to remodel their GABA-ergic synapses onto output neurons.

### Plasmin cleaves EphA4 at R497 in vitro, ex vivo and in vivo in the central amygdala after stress

We next set to identify the molecular substrate targeted by plasmin to regulate the function of GABA-ergic synapses in the CEA. We have previously shown that postsynaptic Eph-receptor tyrosine kinases are subject to cleavage by a different extracellular serine protease (neuropsin) and this process can regulate the activity of excitatory (glutamatergic) synapses in the amygdala (13). To investigate whether plasmin could utilize a similar mechanism to regulate inhibitory synapses, we treated neuroblastoma SH-SY5Y cells with tPA/plasminogen (to generate active plasmin) and studied the cleavage of various Eph receptors by Western blotting. We found that increasing concentrations of plasmin (6-120 nM) caused gradual disappearance of the native EphA4 band, consistent with cleavage of EphA4 (Fig. 2a, n=3-4 per group, p<0.001). The density of bands corresponding to other Eph receptors (B2, B4, B6), their ephrin ligand ephrinB2 or unrelated receptor p75/GFR were not affected by the plasmin treatment (Supplementary Fig. 2). This result indicates that plasmin specifically cleaves EphA4, but not other Eph receptors.

**Figure 2:**
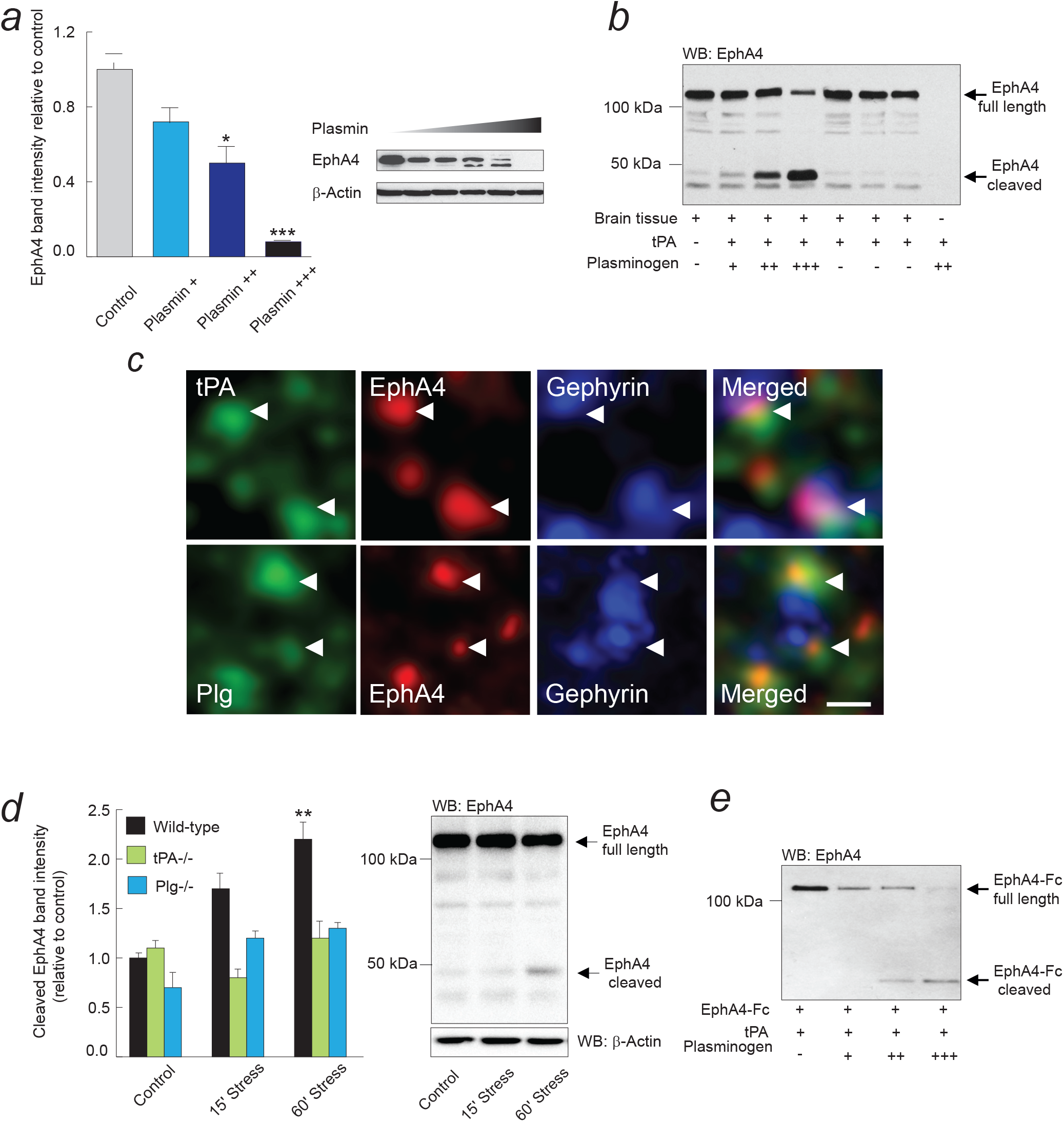
tPA-activated plasmin cleaves EphA4 in vitro and in vivo. **(a)** Treatment of SHSY-5Y cells with increasing concentrations of plasmin (14 nM of tPA and 6, 12 or 120 nM of plasminogen, shown as +, ++ or +++, respectively) reduced the density and eventually caused disappearance of the EphA4 band in Western blotting, consistent with plasmin cleaving EphA4 (F_(3, 12)_= 14.6; p<0.05 and p<0.001). Plasmin cleavage of EphA4 was specific because other Eph receptors remained intact (Supplementary Fig. 2). **(b)** Addition of increasing concentrations of plasmin (as above) to the mouse amygdala homogenate caused a decrease in the density of the native EphA4 band and concomitant appearance of a novel ~50 kDa EphA4 band, demonstrating the capacity of plasmin to cleave EphA4 in the brain milieu. EphA4 co-localise with tPA and plasminogen in gephyrin positive synapses **(c)** in the central amygdala, scale bar is 100nm. **(d)** Mice were subjected to restraint stress, their amygdalae dissected and EphA4 visualized by Western blotting. Two-fold increase in the density of EphA4 cleavage band was observed in stressed wild-type (F _(2, 11)_ = 8.7, p< 0.01, WT stress vs WT no-stress), but not in stressed tPA^−/−^ or Plg^−/−^ animals (F _(2, 11)_ = 0.6 and F _(2, 6)_ = 1.9 respectively, p>0.05). Representative Western blots for tPA^−/−^ and Plg^−/−^ animals are shown in Supplementary Fig. 3 **(e)** Treatment of the purified recombinant EphA4-Fc protein with increasing concentrations of plasmin (as in **a**) caused a decrease in the density of the main EphA4 band and a concomitant appearance of a novel ~50 kDa EphA4 band in the Western blotting, demonstrating that plasmin cleaves EphA4 directly. Similar results were obtained when anti-EphA4 N-terminal (instead of C-terminal) antibodies were used (Supplementary Fig. 3 f). Results are shown as mean ± SEM. *p<0.05, **p<0.01, ***p<0.001

We next examined if plasmin could still efficiently cleave EphA4 in the brain milieu despite the abundance of serine protease inhibitors (17, 18). Western blotting revealed that the addition of increasing doses of tPA/plasminogen to amygdala tissue homogenate resulted in a decrease in the density of the native EphA4 band, and concomitant appearance of a novel band at 50 kDa, corresponding to the cleaved form of EphA4 (Fig. 2b, n=3 per group). In order to examine if EphA4, tPA and plasminogen co-localize at inhibitory synapses in the CEl and thus could readily interact, we performed multi-label immunohistochemistry. It revealed that all three proteins co-localize with gephyrin-positive puncta (a marker of GABA-ergic synapses (23)) in the CEl (Fig. 2c, N=3 mice, n=6-9 sections per group).

In order to determine if EphA4 is cleaved in response to an anxiogenic stimulus we subjected wild-type C57/BL6/J mice to acute restraint stress, dissected their amygdalae and examined the levels of the native and cleaved form of EphA4. Western blotting demonstrated that, upon restraint stress, the density of the band corresponding to the plasmin-cleaved form of EphA4 increased at 60 minutes (Fig. 2d, n=3-4 per group, p<0.01). Consistent with the causal role of tPA/plasmin in cleaving EphA4 after stress, the density of this band did not increase in tPA^−/−^ or plasminogen^−/−^ mice (Fig. 2d, c, n=3-4 per group, p>0.05). Overall, our data (Fig. 1–3a-d) indicate that, in response to an anxiogenic stimulus, plasmin cleaves EphA4 in GABA-ergic synapses downstream of PKCδ^+^ interneurons in the CEl.

In addition to directly cleaving substrates, plasmin can also activate other proteases (such as metalloproteinases) (16), which could potentially also cleave EphA4. Therefore, we next investigated if plasmin directly cleaves EphA4 by treating purified recombinant EphA4 protein with increasing concentrations of tPA/plasminogen. Western blotting revealed a decrease, and eventual disappearance, of the native EphA4 band, accompanied by a simultaneous appearance of the cleaved EphA4 band at 50 kDA (Fig. 2e, n=4-5 per group). This result demonstrates that plasmin is capable of cleaving EphA4 directly, without contribution of other proteases.

We next worked to determine the exact tPA/plasmin cleavage site(s) of EphA4. In order to accomplish this, we employed amino-terminal orientated mass spectrometry of substrates (ATOMS) (24). Recombinant EphA4 protein was treated with increasing concentrations of tPA/plasminogen, the N-termini of the resulting EphA4 fragments labelled with ^13^CD_2_O and subjected to quantitative tandem mass spectrometry (24). ATOMS identified five cleavage sites within the EphA4 extracellular domain (four of which were cleaved by plasmin and three by tPA albeit with ~100-times lower efficiency; Fig. 3a, b, n=3 per protease). Consistent with the known consensus plasmin-cleavage sequences (25), all cleavage sites were at the carboxyl side of an amino acid arginine. We found that arginine at position 497 (R497) was the most sensitive to plasmin’s proteolytic activity. The predicted molecular weight of the EphA4 cleaved at R497 corresponded to the EphA4 fragment detected by us in the mouse amygdala after stress (Fig. 2d). This result demonstrates that plasmin preferentially cleaves EphA4 at R497, located within the extracellular fibronectin type-III domain accessible in the synaptic cleft (19, 26).

**Figure 3:**
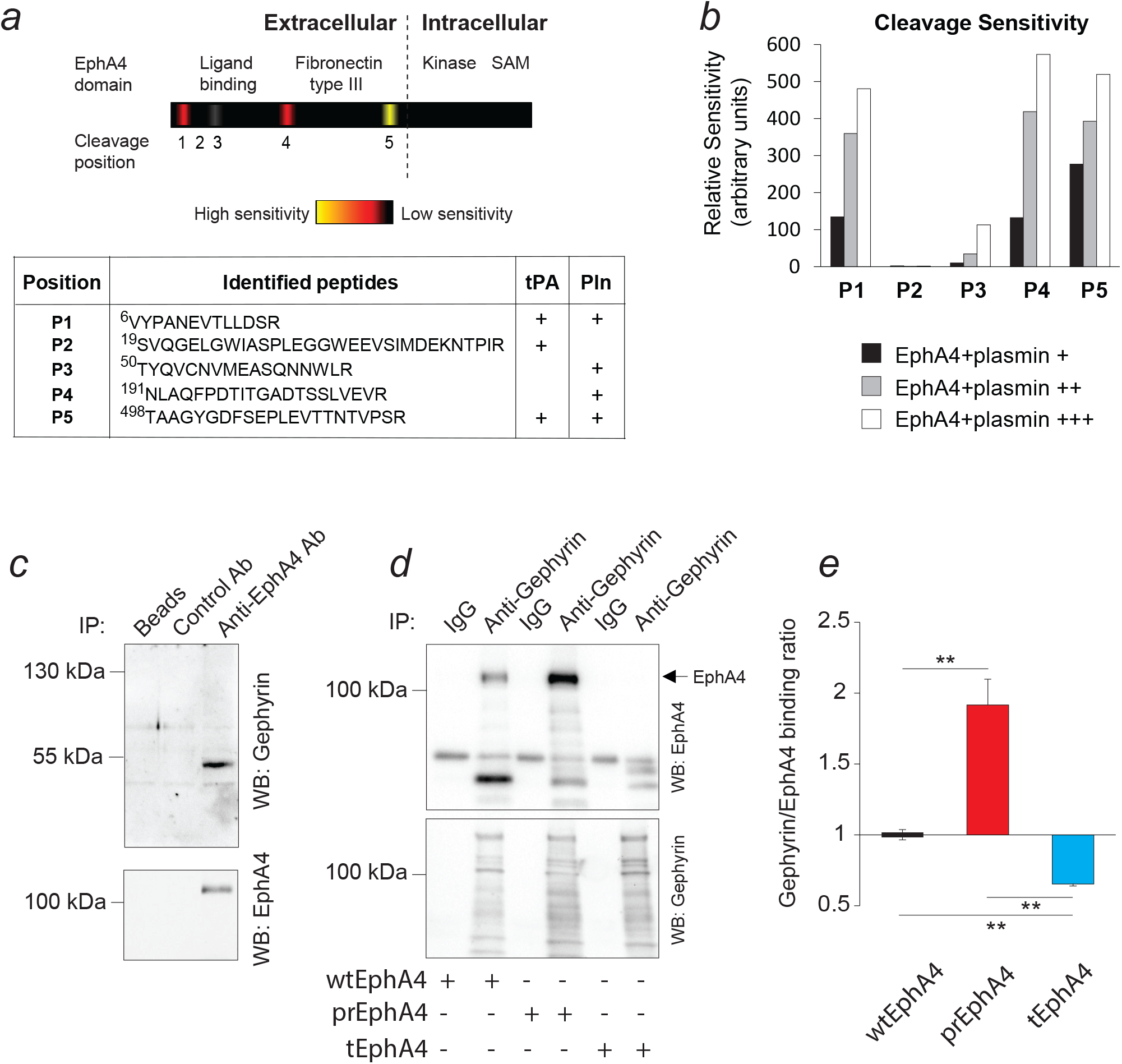
Plasmin cleaves EphA4 at R497 to regulate EphA4/gephyrin binding. **(a)** tPA and plasmin cleavage sites of EphA4 were identified by amino-terminal oriented mass spectrometry of substrates (ATOMS) after treating purified recombinant EphA4-Fc with either tPA or tPA + increasing concentrations of plasminogen (see Methods). Diagram shows the sensitivity and localisation of the cleavage sites relative to the structure of EphA4. Plasmin cleaved EphA4 with 100-fold higher efficiency than tPA. **(b)** The cleavage site P5, located within the fibronectin type-III domain of EphA4, was the most sensitive to cleavage by plasmin. Cleavage at P5 is consistent with the size of the EphA4 plasmin cleavage bands shown in Figure 3 c, d, e. **(c)** Co-immunoprecipitation followed by Western blotting revealed that in the amygdala EphA4 physically interacts with the neuro-skeleton protein gephyrin, a universal regulator of GABA-receptor subunit anchoring. **(d, e)** Further co-immunoprecipitation studies demonstrated that EphA4/gephyrin interaction is dynamic and regulated by plasmin cleavage of EphA4 at R497. EphA4/gephyrin binding is enhanced upon the expression of plasmin-resistant variant of EphA4 (prEphA4), and disrupted upon the expression of EphA4 truncated at R497 (tEphA4) (F_(2, 6)_ = 42.64, P=0.0003, **P<0.01)

### Cleavage of EphA4 at R497 regulates gephyrin binding

Our earlier microarray-based analysis of amygdala signalling pathways pointed towards a GABA-ergic synapse as the main functional unit regulated by the tPA/plasmin system, with the expression of seven GABA-receptor subunits altered by the deletion of the tPA gene (Fig. 1). Because we found EphA4 to be the principal target of the tPA/plasmin system in the amygdala, we hypothesised that, in order to convey the effect of tPA/plasmin on GABA-ergic synapses, EphA4 may physically interact with gephyrin - the master regulator of GABA-receptor subunit anchoring (23, 27). To test this possibility, we immunoprecipitated EphA4 from mouse amygdalae using anti-EphA4 antibody and probed the material with antibodies against gephyrin. Indeed, Western blotting clearly showed that EphA4 physically interacts with gephyrin (Fig. 3c, n=3) consistent with previous immunohistochemistry confirming a high degree of co-localization of EphA4 and gephyrin in the same synaptic terminals (see Fig. 2c).

In order to investigate if plasmin-induced cleavage affects the EphA4/gephyrin interaction we generated a plasmin cleavage-resistant EphA4 variant (prEphA4) through R497Q substitution. We also created an EphA4 variant truncated at R497 (tEphA4) to mimic EphA4 cleavage product generated by plasmin. Analysis of these constructs expressed in a neuroblastoma Neuro-2a cell line confirmed their correct size, resistance to cleavage by plasmin and appropriate cell membrane localisation (Supplementary Fig. 3a-c). We then expressed either the wild-type EphA4 (wtEphA4), prEphA4 or tEphA4 in Neuro-2a cells and performed co-immunoprecipitation studies that confirmed a strong interaction of wtEphA4 with gephyrin (Fig. 3d). Strikingly, EphA4/gephyrin binding was enhanced when prEphA4 was expressed instead of wtEphA4, and disrupted upon the expression of tEphA4 (Fig. 3d-e, n=3 per group p<0.01). Altogether, our results (Fig. 1–4) suggest that EphA4 regulates GABA-receptor subunit synaptic anchoring in a plasmin-dependent manner through its dynamic, physical interaction with gephyrin.

### Cleavage of EphA4 at R497 regulates dendritic spine morphology

Gephyrin is a postsynaptic neuroskeletal protein that, in addition to controlling synaptic anchoring of GABA-receptor subunits, regulates the morphology of postsynaptic terminals and dendritic spines in particular (23, 28). CEl PKCδ^+^ interneurons form axo-dendritic GABA-ergic synaptic contacts with medium spiny neurons in the amygdala (3, 21, 22). This raises the possibility that the morphology of dendritic spines receiving presynaptic boutons from tPA-releasing PKCδ^+^ interneurons could be regulated by the plasmin/EphA4/gephyrin signalling unit. To investigate this hypothesis, we expressed wtEphA4, prEphA4 or tEphA4 together with enhanced green fluorescent protein (EGFP) in primary cultured amygdala neurons and examined their dendritic spine morphology by confocal microscopy. Morphometric analysis revealed that compared to wtEphA4, the expression of prEphA4 promoted the formation of short, mushroom-like spines, while the expression of trEphA4 facilitated the formation of long spines with narrow heads (Fig. 4, N=9-15 neurons per group, >1000 spines per group analysed, p<0.001; Supplementary Fig. 4). Thus, our data suggest that cleavage of EphA4 at R497 by plasmin favours the formation of “thin” dendritic spines.

**Figure 4:**
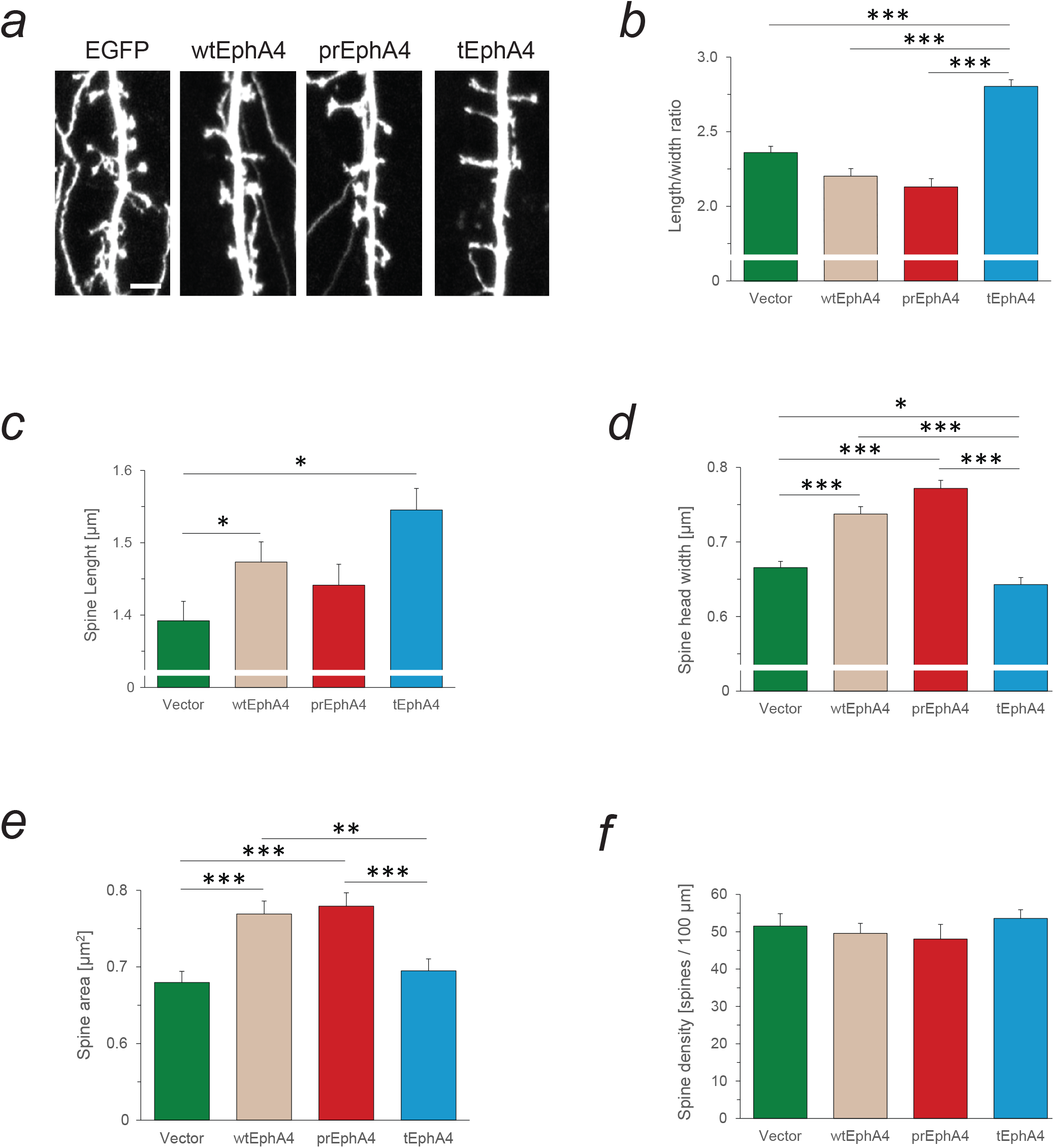
Cleavage of EphA4 at R497 regulates dendritic spine morphology. **(a)** Examples of primary mouse amygdala neurons over-expressing wtEphA4, prEphA4 and tEphA4 variants, scale bar is 3µm. (**b**) Over-expression of prEphA4 favours formation of short dendritic spines with wide heads, while tEphA4 promotes generation of long spines with thin heads (for length/width ratio H(1) = 97.09, P<0.0001) **(c)** Comparison of dendritic spine length. Over-expression of tEphA4 in mouse amygdala neurons resulted in formation of longer spines compared to those transfected with EGFP-containing control vector (H(1) =20.64, P<0.0001) **(d)** Comparison of spine head width. Over-expression of prEphA4 resulted in formation of spines with wide heads, while overexpression of tEphA4 promoted formation of spines with narrow heads (H(1) = 125.3, P<0.0001). **(e)** Comparison of dendritic spine area. Over-expression of either wtEphA4 or prEphA4 led to a formation of larger spines, while overexpression of tEphA4 was devoid of such an effect (H(1) = 33.87, P<0.0001). **(f)** Dendritic spine density was not affected by the overexpression of EphA4 variants (F _(3, 105)_=0.68, P>0.05). The total number of spines analyzed was 1131 (13 neurons) for wtEphA4, 1088 (14 neurons) for prEphA4, 1173 (15 neurons) for tEphA4 and 1241 (9 neurons) for a control construct (containing EGFP only). Results are shown as mean ± SEM. *p<0.05, **p<0.01, ***p<0.001

### Plasmin-mediated cleavage of EphA4 at R497 is critical for the development of stress-induced anxiety

CEl PKCδ^+^ interneurons are critical components of the neural circuit of anxiety (3-5, 21). In particular, recent optogenetic studies support the role of CEl PKCδ^+^ interneurons for control of anxiety through feedforward inhibition of CEm neurons projecting to subcortical motor and autonomic nuclei^4^. In order to investigate whether the tPA/plasmin/EphA4/gephyrin signalling pathway is necessary and/or sufficient for the expression of anxiety we transduced wtEphA4, prEphA4 or tEphA4 (or empty UbC-EGFP vector) into the mouse central amygdala bilaterally using lentiviral gene transfer (Fig. 5a). We then subjected the mice to restraint stress and measured their levels of anxiety using the elevated-plus maze. This test measures the tendency of mice to avoid open, illuminated and elevated arms of the maze, which they perceive as unsafe (29). Open-arm avoidance is more pronounced when the level of anxiety is high (4, 13). Our experiments demonstrated that the EphA4 constructs did not produce a behavioural response in stress-naïve mice, but profoundly (and differentially) altered anxiety profiles in response to a stressful stimulus (Fig. 5 b, c, n=7-19 per group). Consistent with previous reports (13, 14), restraint stress reduced the time control mice (injected with the UbC-EGFP vector) spent exploring the open arms of the maze, consistent with increased anxiety (Fig. 5 b, c, n=18-19 per group, p<0.01). However, mice in which either wtEphA4 or prEphA4 was overexpressed did not show this stress-induced decrease in open-arm exploration, indicative of low levels of anxiety (n=7 per group, p>0.05). In stark contrast, mice in which tEphA4 was overexpressed exhibited dramatically elevated anxiety levels in response to restraint stress (Fig. 5 b, c, n=7 per group, p<0.001). This result demonstrates that plasmin-dependent cleavage of EphA4 at R497 in CEl is necessary for the expression of stress-induced anxiety-like behaviour.

**Figure 5:**
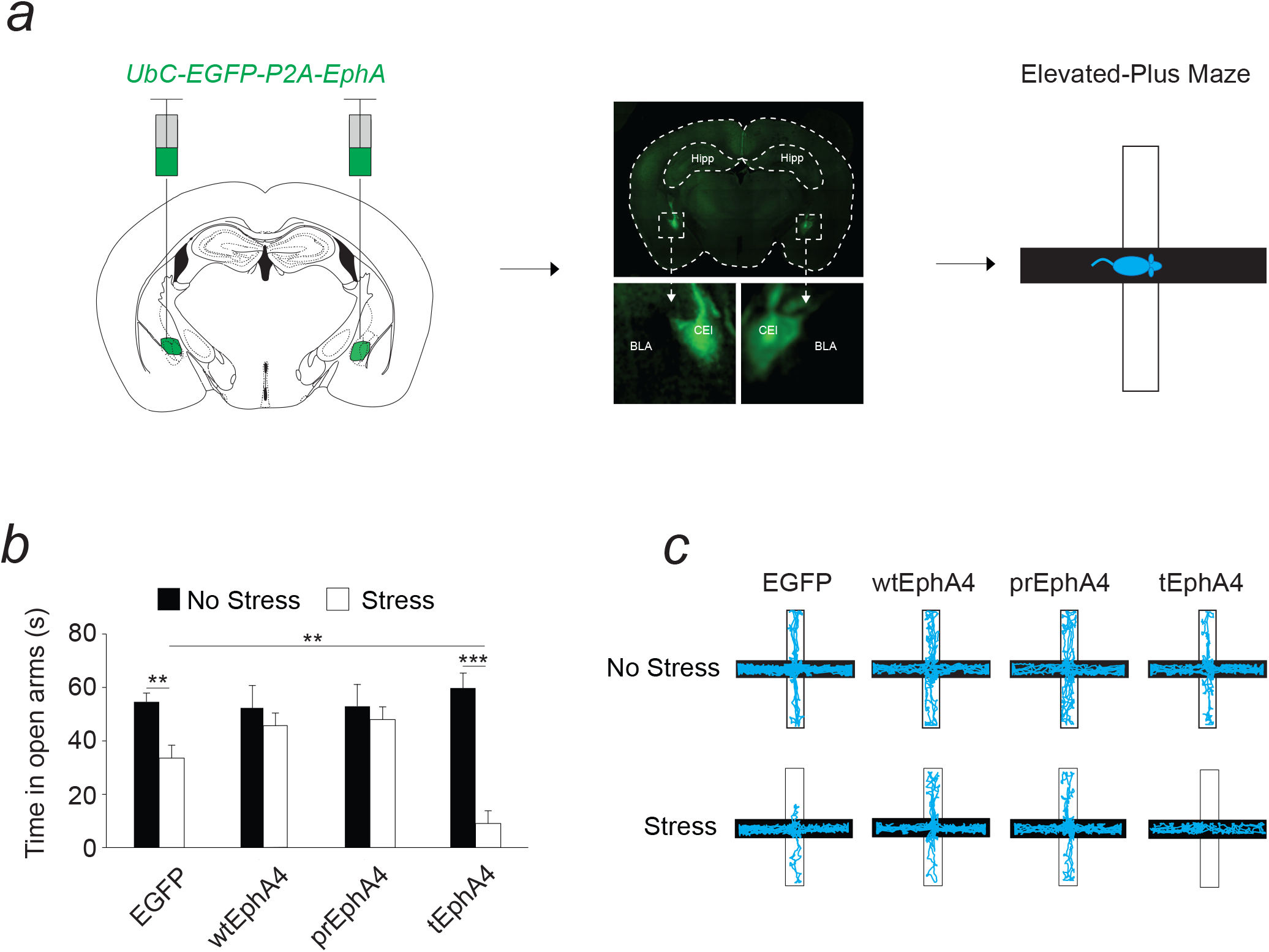
Cleavage of EphA4 at R497 is critical for the development of stress-induced anxiety. **(a)** UbC-EGFP-P2A-EphA lentiviral vectors, containing either wtEphA4, prEphA4 or tEphA4 (or EGFP as control), were bilaterally injected into the central amygdala (CEA) and mice were allowed to recover for three weeks to allow the transgene expression. Then, mice were subjected to restraint stress (or left undisturbed) and their anxiety levels measured in the elevated-plus maze. Representative injection sites, identified by the expression of EGFP, are shown in the middle panel. **(b)** In UbC-EGFP-injected mice, restraint stress caused a decrease in the time spent in open arms, indicative of elevated levels of anxiety. Overexpression of either wtEphA4 or prEphA4 in the same vector prevented the development of stress-induced anxiety, supporting the view that uncleaved EphA4 promotes anxiolysis. Consistently, overexpression of tEphA4 in CEA dramatically enhanced stress-induced anxiety, altogether demonstrating that plasmin cleavage of EphA4 at R497 is necessary for this process (F_(7, 65)_ = 6.796; P<0.0001; **p<0.01, ***p<0.01). **(c)** Representative elevated-plus maze traces of mice subjected to restraint stress. Results are shown as mean ± SEM. **p<0.01, ***p<0.001

## Discussion

Here we report a new signalling cascade that bi-directionally controls activity of central amygdala interneurons and consequently stress-induced anxiety-like behaviour. The above pathway is initiated by tPA-mediated activation of plasminogen, resulting in plasmin-dependent cleavage of the EphA4 receptor. Cleavage of EphA4 leads to its dissociation from the postsynaptic scaffold protein gephyrin, and culminates in structural and functional rearrangements of GABA-ergic synapses to regulate anxiety.

### Regulation of inhibitory GABA-ergic amygdala synapses by tPA and plasmin

In order to determine the mechanism of action of tPA in the amygdala we used an unbiased approach to identify the structural or functional amygdala elements affected by the deletion of this protease. Our microarray study demonstrated that tPA predominantly controls GABA-ergic amygdala synapses. This finding is consistent with tPA localization in the primarily inhibitory region of the amygdala (CEl) and its selective expression by GABA-ergic PKCδ^+^ interneurons. In particular, we found that tPA regulates GABA-receptor subunit composition of inhibitory synapses in CEl. This unexpected finding contrasts with the previously reported role of the tPA/plasmin system in excitatory signalling in the cortex (30), and suggests that the mechanisms the tPA/plasmin system uses to exert its effects are region-dependent. Although we cannot entirely exclude the possibility that tPA, when released from inhibitory terminals into the synaptic space, could potentially diffuse into adjacent glutamatergic synapses to modulate excitatory signalling, we did not find evidence of such a mechanism in our study. Quite the opposite, our microarray, immunohistochemistry and in situ hybridization data collectively point to the inhibitory signalling as a sole *modus operandi* of the tPA/plasmin system in CEl.

### Cellular origins of tPA and plasminogen in the amygdala

Previously reported immunohistochemistry data reproducibly showed a diffuse extracellular pattern of tPA expression (14, 31), consistent with its release into the brain parenchyma (14, 15), but precluding a meaningful identification of its source. Potentially, tPA could be synthesized outside the amygdala, transported alongside axonal projections and deposited in the central amygdala nucleus. However, high levels of tPA mRNA within the amygdala detected by us make this possibility unlikely. Instead, we found that tPA is both produced and released locally by the central amygdala interneurons. In particular, we found the abundance of tPA mRNA in a specific subclass of GABA-ergic interneurons located within CEl, characterized by the expression of protein kinase Cδ (PKCδ^+^ cells). Our findings are consistent with previously published low-resolution *in situ* hybridisation data, showing a diffuse tPA mRNA signal in the amygdala of mice (32). Thus, our study demonstrates that amygdala tPA originates from inhibitory PKCδ^+^ neurons and is therefore likely to regulate the activity of their inhibitory synapses onto CEm cells.

Previous attempts to demonstrate the presence of plasminogen in the healthy mouse brain have been partially successful, but hampered by low plasminogen mRNA levels in the central nervous system and low sensitivity of methods utilized for protein detection (31, 33). Using high-sensitivity immunohistochemistry, we demonstrated that plasminogen, albeit at lower levels than tPA, is expressed in the central nucleus of the amygdala. Therefore, both components of the tPA/plasmin system are expressed in the above brain structure to form a functional proteolytic cascade.

### Cleavage of Eph receptors by plasmin in synaptic physiology

We next set to identify the substrate that tPA and/or plasmin cleave in the amygdala to modulate GABA-ergic signalling. We found that plasmin (upon activation by tPA) cleaves the membrane tyrosine kinase receptor EphA4, a member of the ephrin receptor family highly expressed in the brain (34). EphA4 is known to play a role in amygdala physiology (35), undergo proteolytic cleavage (19), and to control various forms of experience-driven neuronal plasticity (34). Importantly, our study is the first to identify an EphA4 cleavage site, attribute it to a specific protease and demonstrate its role in inhibitory plasticity. The cleavage site identified by us is located within the fibronectin type III domain of EphA4 and is distinct from previously reported EphA4 cleavage (19).

Based on our current and previous findings we hypothesize that cleavage of different Eph receptors by distinct serine proteases serves as a universal mechanism of controlling both the excitatory and inhibitory plasticity in the amygdala (13–15). We found previously that another serine protease, neuropsin, is highly expressed in the excitatory basolateral amygdala, where it cleaves a member of the ephrin receptor family, EphB2. Neuropsin-mediated cleavage of EphB2 controls excitatory signalling and anxiety-like behaviour by regulating EphB2/NMDA receptor binding (13). In the present study we found that plasmin-mediated cleavage of EphA4 in the central amygdala controls inhibitory, GABA-ergic signalling. Interestingly, both neuropsin (13) and plasmin (this study) cleave Eph receptors within their fibronectin type III domains. Therefore, it appears that an orchestrated action of different serine proteases (neuropsin vs. tPA/plasmin) in the amygdala regions characterized by distinct neurotransmitter profiles (the excitatory basolateral vs. the inhibitory central amygdala) can proteolytically modulate different members of the Eph receptor family (EphB2 vs. EphA4), thus controlling the excitatory / inhibitory balance at the amygdala network level, and consequently anxiety-like behaviour.

### EphA4 interaction with gephyrin and dendritic spine plasticity

Unexpectedly, we found that EphA4 directly binds to gephyrin and that EphA4/gephyrin interaction is regulated by the tPA/plasmin system. Gephyrin is located in dendritic spines of inhibitory neurons and converges extracellular signals in order to translate them into adjustments in the strength of GABA-ergic transmission (23, 36). This can be achieved by altering the receptor-anchoring properties of the gephyrin-associated molecular machinery, thus changing the GABA-receptor subunit composition at inhibitory synapses over short timescales (23, 36). Although we did not directly measure the fluctuations in GABA-ergic transmission in response to EphA4 cleavage, the results of our microarray study clearly demonstrate tPA-dependent alterations in the GABA-receptor subunit composition in the amygdala. Such alterations have previously been shown to modulate inhibitory transmission (37). Thus, the tPA/plasmin/EphA4/gephyrin pathway promotes adaptive forms of inhibitory plasticity in CEl.

Although the presence of dendritic spines is a typical attribute of excitatory neurons, a number of histological studies have demonstrated that a subset of GABA-ergic neurons in the amygdala carry dendritic spines (38, 39). Interestingly, dendritic spines are often co-innervated both by the excitatory and inhibitory inputs (40–43). Because each dendritic spine typically bears only a few synapses, additional inhibitory input is uniquely poised to override the impact of the excitatory transmission on the spine physiology, which may result in substantial, GABA-dependent re-arrangements of dendritic spine morphology (40–42, 44). We, therefore, set to examine if the plasmin/EphA4/gephyrin pathway controls dendritic spine geometry. The morphology of interneuronal dendritic spines follows a typical pattern observed in excitatory pyramidal neurons, with mushroom, stubby, thin, and filopodia-like structures (43). We found that the expression of the plasmin-truncated variant of EphA4 causes an increase in the proportion of immature dendritic protrusions (filopodia) while decreasing the percentage of the mature, mushroom-like spines in the amygdala neurons. Gephyrin has previously been reported to regulate dendritic spine maturation through α5-GABA-A receptor signalling (28), but the abundant literature demonstrating the involvement of EphA4 in dendritic spine structural plasticity has not to date implicated gephyrin in these processes (45–48). Our study describes a new mechanism of controlling dendritic spine morphology through plasmin-mediated cleavage of EphA4 which in turn triggers gephyrin-dependent alterations of synaptic GABA-receptor subunit expression profiles. These re-adjustments of the inhibitory synapse composition either offset or facilitate the impact of the excitatory transmission on spine morphology to bi-directionally regulate anxiety-like behaviour.

## Conclusions

In summary, we identified a novel trans-synaptic signalling cascade that regulates the function of CEl PKCδ^+^ GABA-ergic synapses - a key component of the amygdala circuit of anxiety. Our studies favour a model (Supplementary Fig. 5) where, in response to stress, CEl PKCδ^+^ interneurons release tPA at their GABA-ergic synapses. Subsequently, tPA converts inactive plasminogen to active plasmin which cleaves postsynaptic EphA4 at R497. Plasmin-mediated cleavage triggers dissociation of EphA4 from its intracellular partner, gephyrin - a neuroskeletal GABA-receptor subunit anchoring protein. Dynamic EphA4/gephyrin interaction leads to remodelling of the postsynaptic terminal, promoting the formation of thin dendritic spines and altering the amygdala GABA-receptor subunit expression profile. The above rearrangements support the development of stress-induced anxiety. Molecular manipulation of the tPA/plasmin/EphA4/gephyrin signalling cascade can either switch on or switch off the development of stress-induced anxiety in mice, and may thus provide a novel therapeutic avenue to increase stress resilience and treat anxiety disorders in humans.

## Acknowledgements

The study was supported by the Marie Sklodowska Curie ITN grant “Extrabrain” and ECMNet COST Action (European Commission) to R.P and L.K, The Leverhulme Project Grant to R.P, Cleopatra Trust grant to R.P., National Science Centre grant (UMO-2015/17/B/NZ3/00557) to J.W. M.M. and I.F., Foundation for Polish Science TEAM grant (Team/2016-1/6) to L.K. and Polish National Science Centre Grant number 2013/08/A/NZ3/00848 to R. Przewlocki. We would like to thank Megan Jackson for her involvement in the immunohistochemistry experiments.

## Conflict of interest

The authors declare that they have no conflict of interest.

Supplementary information is available at MP’s website.

## References

1. McEwen BS, Bowles NP, Gray JD, Hill MN, Hunter RG, Karatsoreos IN, et al. Mechanisms of stress in the brain. Nat Neurosci. 2015;18(10):1353–63.

2. Ehrlich I, Humeau Y, Grenier F, Ciocchi S, Herry C, Luthi A. Amygdala inhibitory circuits and the control of fear memory. Neuron. 2009;62(6):757–71.

3. Janak PH, Tye KM. From circuits to behaviour in the amygdala. Nature. 2015;517(7534):284–92.

4. Botta P, Demmou L, Kasugai Y, Markovic M, Xu C, Fadok JP, et al. Regulating anxiety with extrasynaptic inhibition. Nat Neurosci. 2015;18(10):1493–500.

5. Haubensak W, Kunwar PS, Cai H, Ciocchi S, Wall NR, Ponnusamy R, et al. Genetic dissection of an amygdala microcircuit that gates conditioned fear. Nature. 2010;468(7321):270–6.

6. Sandi C, Haller J. Stress and the social brain: behavioural effects and neurobiological mechanisms. Nat Rev Neurosci. 2015;16(5):290–304.

7. Bandelow B, Michaelis S. Epidemiology of anxiety disorders in the 21st century. Dialogues Clin Neurosci. 2015;17(3):327–35.

8. Kessler RC, Ruscio AM, Shear K, Wittchen HU. Epidemiology of anxiety disorders. Curr Top Behav Neurosci. 2010;2:21–35.

9. Tovote P, Fadok JP, Luthi A. Neuronal circuits for fear and anxiety. Nat Rev Neurosci. 2015;16(6):317–31.

10. Roozendaal B, McEwen BS, Chattarji S. Stress, memory and the amygdala. Nat Rev Neurosci. 2009;10(6):423–33.

11. Capogna M. GABAergic cell type diversity in the basolateral amygdala. Curr Opin Neurobiol. 2014;26:110–6.

12. Cai H, Haubensak W, Anthony TE, Anderson DJ. Central amygdala PKC-delta(+) neurons mediate the influence of multiple anorexigenic signals. Nat Neurosci. 2014;17(9):1240–8.

13. Attwood BK, Bourgognon JM, Patel S, Mucha M, Schiavon E, Skrzypiec AE, et al. Neuropsin cleaves EphB2 in the amygdala to control anxiety. Nature. 2011;473(7347):372–5.

14. Pawlak R, Magarinos AM, Melchor J, McEwen B, Strickland S. Tissue plasminogen activator in the amygdala is critical for stress-induced anxiety-like behavior. Nat Neurosci. 2003;6(2):168–74.

15. Pawlak R, Rao BS, Melchor JP, Chattarji S, McEwen B, Strickland S. Tissue plasminogen activator and plasminogen mediate stress-induced decline of neuronal and cognitive functions in the mouse hippocampus. Proc Natl Acad Sci U S A. 2005;102(50):18201–6.

16. Tsilibary E, Tzinia A, Radenovic L, Stamenkovic V, Lebitko T, Mucha M, et al. Neural ECM proteases in learning and synaptic plasticity. Prog Brain Res. 2014;214:135–57.

17. Melchor JP, Strickland S. Tissue plasminogen activator in central nervous system physiology and pathology. Thromb Haemost. 2005;93(4):655–60.

18. Samson AL, Medcalf RL. Tissue-type plasminogen activator: a multifaceted modulator of neurotransmission and synaptic plasticity. Neuron. 2006;50(5):673–8.

19. Gatto G, Morales D, Kania A, Klein R. EphA4 receptor shedding regulates spinal motor axon guidance. Curr Biol. 2014;24(20):2355–65.

20. Matys T, Pawlak R, Matys E, Pavlides C, McEwen BS, Strickland S. Tissue plasminogen activator promotes the effects of corticotropin-releasing factor on the amygdala and anxiety-like behavior. Proc Natl Acad Sci U S A. 2004;101(46):16345–50.

21. Hunt S, Sun Y, Kucukdereli H, Klein R, Sah P. Intrinsic Circuits in the Lateral Central Amygdala. eNeuro. 2017;4(1).

22. Krabbe S GJ, Lüthi A. Amygdala Inhibitory Circuits Regulate Associative Fear Conditioning. Biological Psychiatry. 2017;in press(https://doi.org/10.1016/j.biopsych.2017.10.006).

23. Tyagarajan SK, Fritschy JM. Gephyrin: a master regulator of neuronal function? Nat Rev Neurosci. 2014;15(3):141–56.

24. Doucet A, Overall CM. Amino-Terminal Oriented Mass Spectrometry of Substrates (ATOMS) N-terminal sequencing of proteins and proteolytic cleavage sites by quantitative mass spectrometry. Methods Enzymol. 2011;501:275–93.

25. Troll W, Sherry S, Wachman J. The action of plasmin on synthetic substrates. J Biol Chem. 1954;208(1):85–93.

26. Kania A, Klein R. Mechanisms of ephrin-Eph signalling in development, physiology and disease. Nat Rev Mol Cell Biol. 2016;17(4):240–56.

27. Beuter S, Ardi Z, Horovitz O, Wuchter J, Keller S, Saha R, et al. Receptor tyrosine kinase EphA7 is required for interneuron connectivity at specific subcellular compartments of granule cells. Sci Rep. 2016;6:29710.

28. Brady ML, Jacob TC. Synaptic localization of alpha5 GABA (A) receptors via gephyrin interaction regulates dendritic outgrowth and spine maturation. Dev Neurobiol. 2015;75(11):1241–51.

29. Kumar V, Bhat ZA, Kumar D. Animal models of anxiety: a comprehensive review. J Pharmacol Toxicol Methods. 2013;68(2):175–83.

30. Nicole O, Docagne F, Ali C, Margaill I, Carmeliet P, MacKenzie ET, et al. The proteolytic activity of tissue-plasminogen activator enhances NMDA receptor-mediated signaling. Nat Med. 2001;7(1):59–64.

31. Salles FJ, Strickland S. Localization and regulation of the tissue plasminogen activator-plasmin system in the hippocampus. J Neurosci. 2002;22(6):2125–34.

32. Sappino AP, Madani R, Huarte J, Belin D, Kiss JZ, Wohlwend A, et al. Extracellular proteolysis in the adult murine brain. J Clin Invest. 1993;92(2):679–85.

33. Taniguchi Y, Inoue N, Morita S, Nikaido Y, Nakashima T, Nagai N, et al. Localization of plasminogen in mouse hippocampus, cerebral cortex, and hypothalamus. Cell Tissue Res. 2011;343(2):303–17.

34. Klein R. Bidirectional modulation of synaptic functions by Eph/ephrin signaling. Nat Neurosci. 2009;12(1):15–20.

35. Deininger K, Eder M, Kramer ER, Zieglgansberger W, Dodt HU, Dornmair K, et al. The Rab5 guanylate exchange factor Rin1 regulates endocytosis of the EphA4 receptor in mature excitatory neurons. Proc Natl Acad Sci U S A. 2008;105(34):12539–44.

36. Choii G, Ko J. Gephyrin: a central GABAergic synapse organizer. Exp Mol Med. 2015;47:e158.

37. Barberis A, Bacci A. Plasticity of GABAergic synapses. Front Cell Neurosci. 2015:1–175.

38. Cassell MD, Gray TS, Kiss JZ. Neuronal architecture in the rat central nucleus of the amygdala: a cytological, hodological, and immunocytochemical study. J Comp Neurol. 1986;246(4):478–99.

39. McDonald AJ. Cytoarchitecture of the central amygdaloid nucleus of the rat. J Comp Neurol. 1982;208(4):401–18.

40. Jasinska M, Siucinska E, Glazewski S, Pyza E, Kossut M. Characterization and plasticity of the double synapse spines in the barrel cortex of the mouse. Acta Neurobiol Exp (Wars). 2006;66(2):99–104.

41. Knott GW, Quairiaux C, Genoud C, Welker E. Formation of dendritic spines with GABAergic synapses induced by whisker stimulation in adult mice. Neuron. 2002;34(2):265–73.

42. Popov VI, Stewart MG. Complexity of contacts between synaptic boutons and dendritic spines in adult rat hippocampus: three-dimensional reconstructions from serial ultrathin sections in vivo. Synapse. 2009;63(5):369–77.

43. Scheuss V, Bonhoeffer T. Function of dendritic spines on hippocampal inhibitory neurons. Cereb Cortex. 2014;24(12):3142–53.

44. Hering H, Sheng M. Dendritic spines: structure, dynamics and regulation. Nat Rev Neurosci. 2001;2(12):880–8.

45. Attwood BK, Patel S, Pawlak R. Ephs and ephrins: emerging therapeutic targets in neuropathology. Int J Biochem Cell Biol. 2012;44(4):578–81.

46. Clifford MA, Kanwal JK, Dzakpasu R, Donoghue MJ. EphA4 expression promotes network activity and spine maturation in cortical neuronal cultures. Neural Dev. 2011;6:21.

47. Fu WY, Chen Y, Sahin M, Zhao XS, Shi L, Bikoff JB, et al. Cdk5 regulates EphA4-mediated dendritic spine retraction through an ephexin1-dependent mechanism. Nat Neurosci. 2007;10(1):67–76.

48. Murai KK, Nguyen LN, Irie F, Yamaguchi Y, Pasquale EB. Control of hippocampal dendritic spine morphology through ephrin-A3/EphA4 signaling. Nat Neurosci. 2003;6(2):153–60.

