## Supplementary Figures for "Plasmin-mediated cleavage of EphA4 at central amygdala inhibitory synapses controls anxiety"

Supplementary Fig. 1

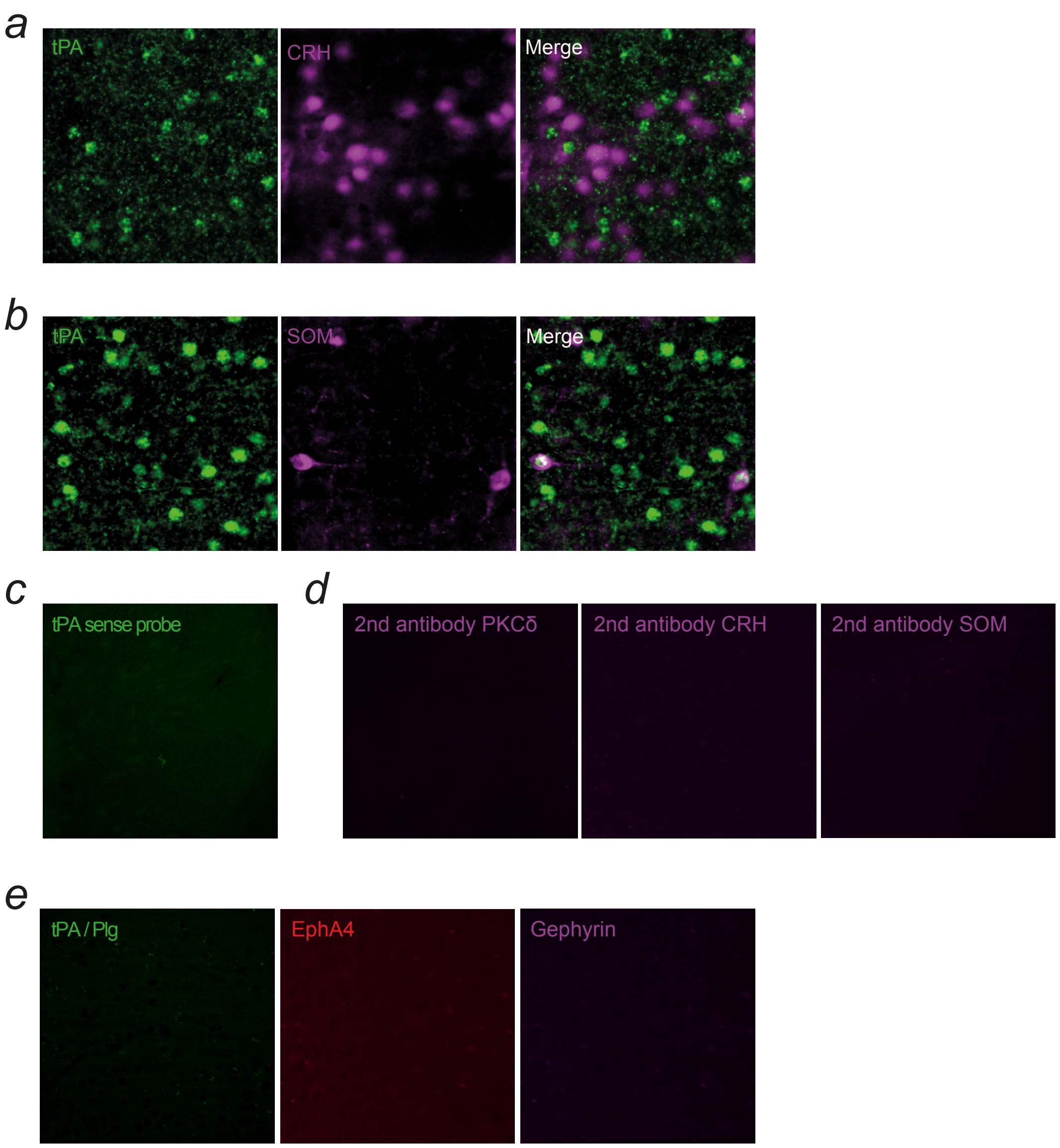

Supplementary Figure 2

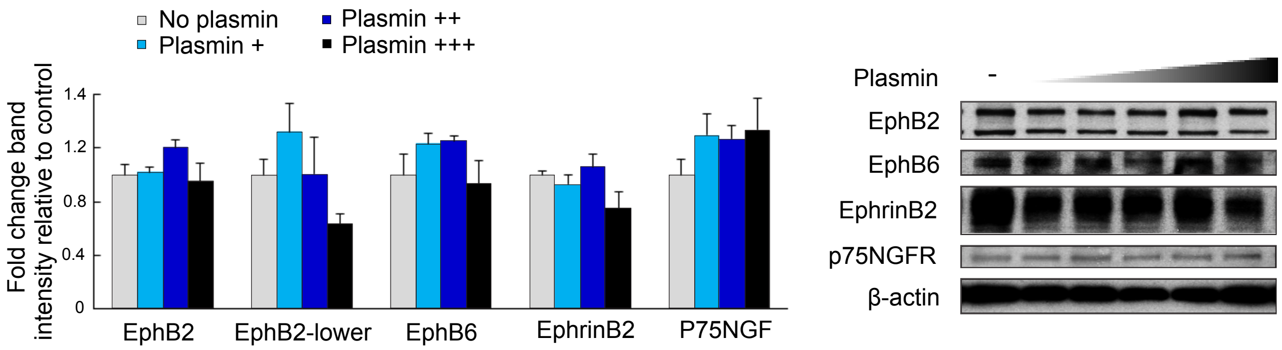

Supplementary Figure 3

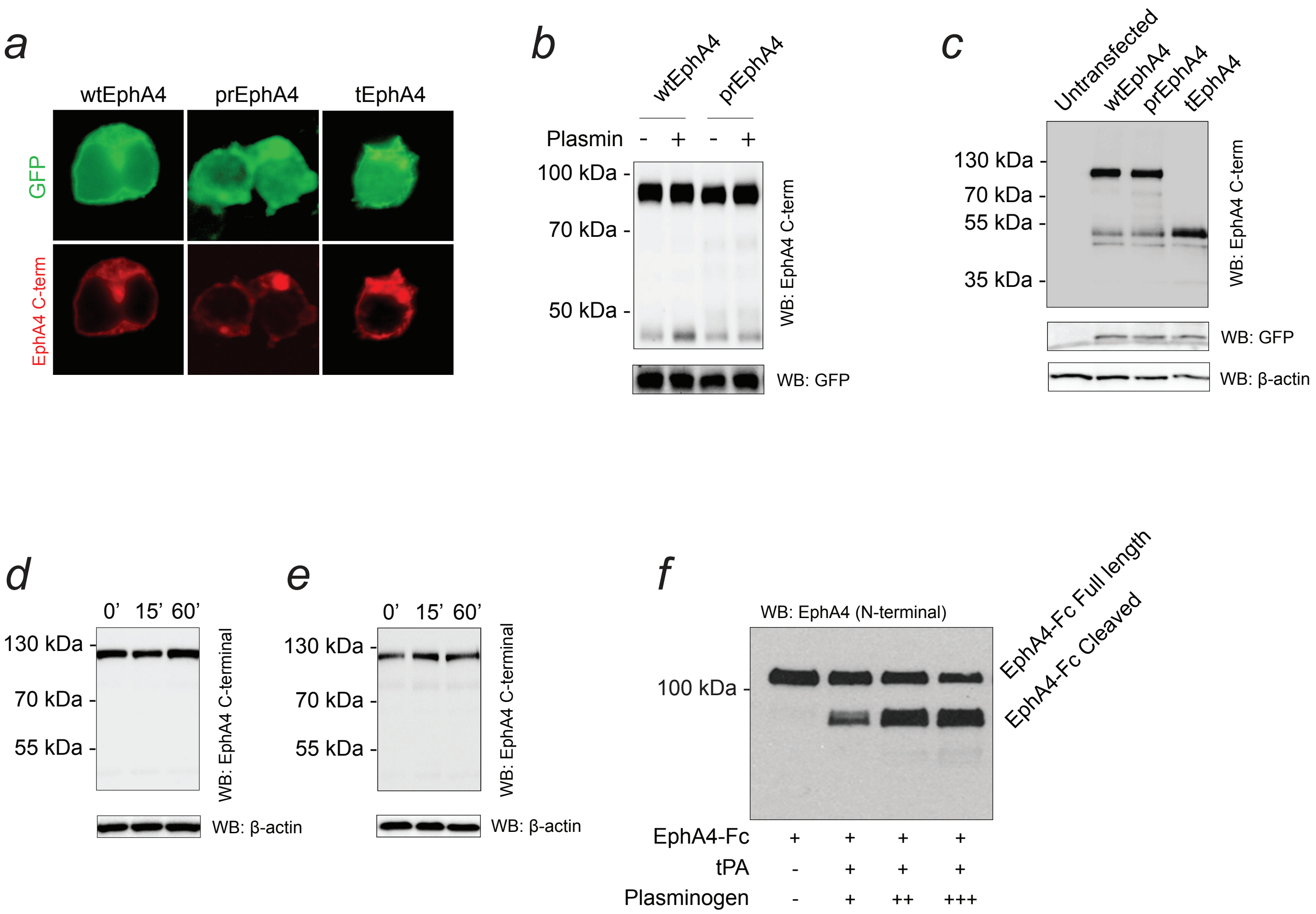

Supplementary Figure 4

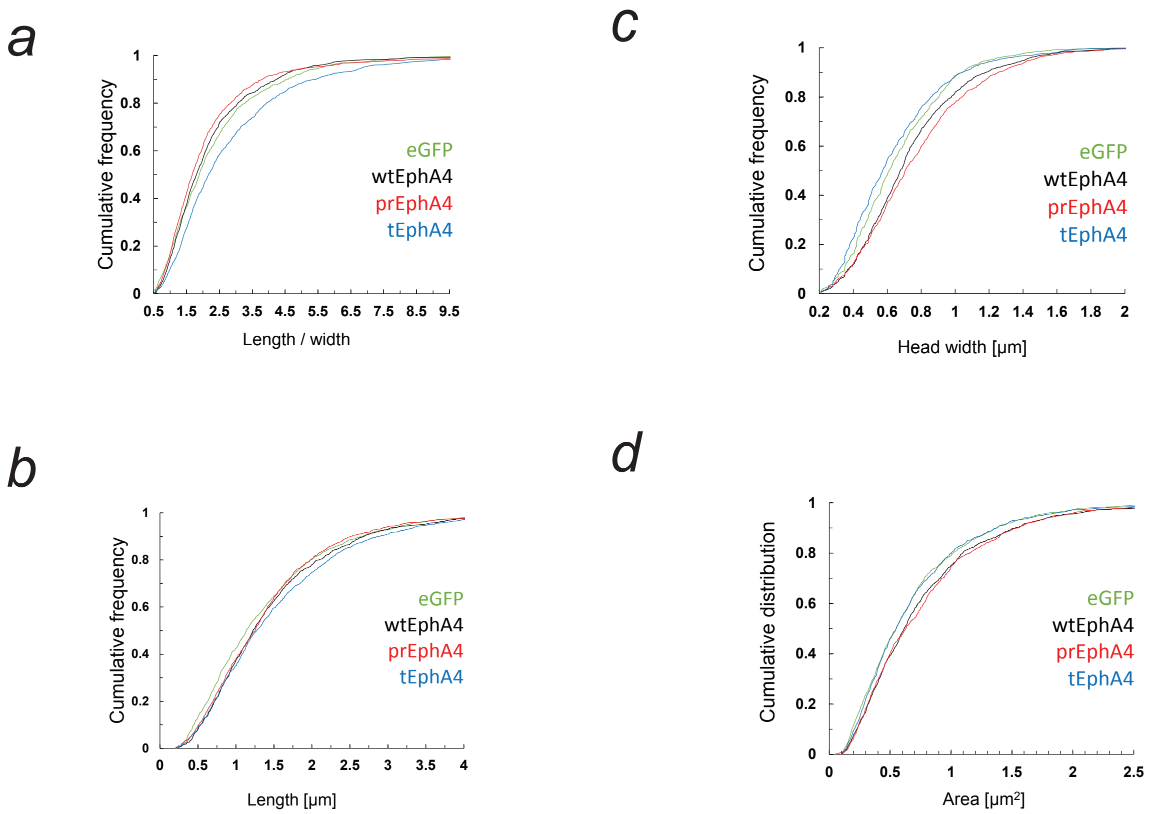

Suppl. Fig. 5

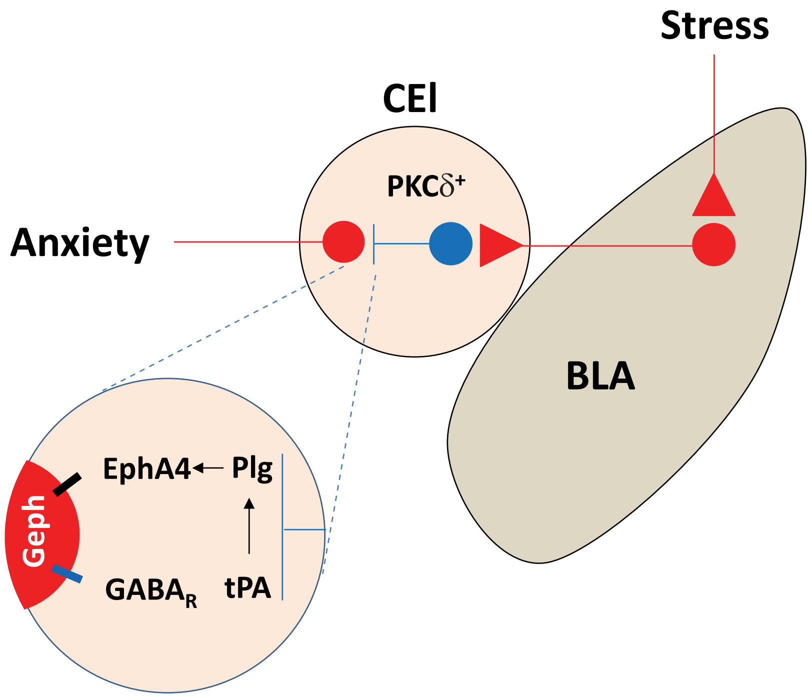

### Supplementary Figure legends:

**Supplementary Figure 1: Specificity of expression of the tPA / plasmin / EphA4 / gephyrin system components in CEl.**

**(a, b)** Fluorescent *in situ* hybridization for the tPA mRNA (green) performed in conjunction with immunohistochemistry for markers of interneuron sub-populations (magenta) revealed that only a small proportion of corticotropin-releasing hormone-positive ($CRH^+$) or somatostatin-positive ($SOM^+$) cells in CEl contained tPA mRNA (p<0.0001 vs $PKC\delta^+$ cells, quantification shown in Figure 1). **(c)** *In situ* hybridization using the tPA mRNA sense probe showing low background fluorescence **(d)** Control immunohistochemistry in which the primary antibodies against PKCδ, CRH and SOM were omitted. **(e)** Endogenous peroxidase inactivation control for tPA and Plg immunohistochemistry and control immunohistochemistry in which the primary antibody against EphA4 or gephyrin were omitted.

**Supplementary Figure 2: Specificity of the EphA4 cleavage by tPA/plasmin**

SH-SY5Y cells were treated with14 nM of tPA and 6, 12 or 120 nM of plasminogen (shown as +, ++ and +++, respectively) to generate active plasmin, and cleavage of EphB2 (upper and lower band), EphB6, ephrinB2 or p75/GFR cleavage was studied by Western blotting (n=3-4 per group). The results were normalized to β-actin. None of the above proteins were cleaved by plasmin (EphB2 $F_{(3, 12)} = 1.70$; P>0.05, EphB2 lower $F_{(3, 12)} = 2.26$; P>0.05, EphB6 $F_{(3, 12)} = 1.77$; P>0.05;, ephrinB2 $F_{(3, 12)} = 2.40$; P>0.05 and p75/NGF $F_{(3, 12)} = 0.89$; P>0.05), confirming that the cleavage of EphA4 was specific. Results are shown as mean ± SEM.

**Supplementary Figure 3: Verification of the membrane localisation, cleavage resistance and molecular mass of plasmin cleavage-related EphA4 variants**

**(a)** Expression of the plasmin cleavage-related EphA4 variants (wild-type EphA4, plasmin cleavage resistant prEphA4 and truncated tEphA4) in Neuro-2a cells demonstrates their cell membrane localization. Transfection was confirmed by co-expression of EGFP. **(b)** prEphA4 is resistant to cleavage by plasmin. Neuro-2a cells transfected with either wtEphA4 or prEphA4 were treated with tPA + plasminogen and EphA4 analysed by Western blotting. Plasmin caused an increase in the density of the EphA4 cleaved band in wtEphA4- but not in prEphA4-transfected cells. Equal level of transfection between groups was confirmed by the expression of EGFP. **(c)** Characterization of wtEphA4, prEphA4 and tEphA4 variants by Western blotting in Neuro-2a cells. R497Q substitution in prEphA4 did not affect the size or level of expression compared with wtEphA4. Deletion of the EphA4 ectodomain at R497 in tEphA4 produced a receptor whose size corresponded to the plasmin-cleaved (Figure 3 b,e) and the stress-induced (Figure 3d) EphA4 band. Transfection was confirmed by co-expression of EGFP and loading by the expression of β-actin.

**Supplementary Fig. 4: Cleavage of EphA4 at R497 controls dendritic spine morphology.**

Cumulative plot of the spine length/width ratio **(a)**, spine length **(b)**, head width **(c)** and cumulative distribution of spine area **(d)** as shown in Figure 3.

**Supplementary Fig. 5: Control of anxiety in central amygdala synapses by the tPA / plasmin / EphA4 / gephyrin signalling pathway.**

Diagram showing the mechanism of molecular control of anxiety in the central amygdala GABA-ergic synapses. Upon stress, tPA is released into synapses downstream of CEl PKC$\delta^+$ interneurons to activate plasminogen to plasmin. Subsequently, plasmin cleaves EphA4 at R497 to trigger its dissociation from the postsynaptic GABA-receptor subunit anchoring protein, gephyrin. Dynamic EphA4/gephyrin interaction triggers remodelling of the postsynaptic terminal, resulting in the formation of thin dendritic spines with altered GABA-receptor subunit expression profiles. The above synaptic changes facilitate the development of stress-induced anxiety. Excitatory neurons are shown in red, inhibitory interneurons in blue. CEl – lateral division of the central amygdala, BLA – basolateral amygdala, Geph – gephyrin.
